# False gene and chromosome losses affected by assembly and sequence errors

**DOI:** 10.1101/2021.04.09.438906

**Authors:** Juwan Kim, Chul Lee, Byung June Ko, DongAhn Yoo, Sohyoung Won, Adam Phillippy, Olivier Fedrigo, Guojie Zhang, Kerstin Howe, Jonathan Wood, Richard Durbin, Giulio Formenti, Samara Brown, Lindsey Cantin, Claudio V. Mello, Seoae Cho, Arang Rhie, Heebal Kim, Erich D. Jarvis

**Author notes:** Corresponding authors: HBK, EDJ. Both authors contributed equally to this work (Juwan Kim, Chul Lee).

## Abstract

Many genome assemblies have been found to be incomplete and contain misassemblies. The Vertebrate Genomes Project (VGP) has been producing assemblies with an emphasis on being as complete and error-free as possible, utilizing long reads, long-range scaffolding data, new assembly algorithms, and manual curation. Here we evaluate these new vertebrate genome assemblies relative to the previous references for the same species, including a mammal (platypus), two birds (zebra finch, Anna’s hummingbird), and a fish (climbing perch). We found that 3 to 11% of genomic sequence was entirely missing in the previous reference assemblies, which included nearly entire GC-rich and repeat-rich microchromosomes with high gene density. Genome-wide, between 25 to 60% of the genes were either completely or partially missing in the previous assemblies, and this was in part due to a bias in GC-rich 5’-proximal promoters and 5’ exon regions. Our findings reveal novel regulatory landscapes and protein coding sequences that have been greatly underestimated in previous assemblies and are now present in the VGP assemblies.

## Introduction

A multitude of reference genome assemblies of diverse vertebrates have been reported in the literature. However, their completeness and accuracy varies^1–3^. Except for human and a few model species, many of these assemblies have been generated using cost-effective platforms with short reads that are a few hundred (<200 bp, mainly Illumina reads) to a thousand bp (~1 kbp, Sanger reads) in length^4,5^. More recently, the development of novel long-read sequencing platforms (e.g. Pacific Biosciences and Oxford Nanopore), novel long-range scaffolding data (e.g. Bionano optical maps and Hi-C), and novel algorithms, allow the generation of more complete and accurate higher quality assemblies. Utilizing and further developing these technologies, the Vertebrate Genomes Project (VGP) aims to generate as complete and error-free genome assemblies as possible, of all extant vertebrate species^6,7^. The Phase 1 pipeline developed by the VGP is to generate haplotype-phased contigs with PacBio continuous long-reads^8^ (CLR) and scaffold the contigs with linked-reads^9^ (10X Genomics), optical maps^10^ (Bionano Genomics), and Hi-C reads^11^ (Arima Genomics). The resulting assemblies have one of the highest contiguity metrics for vertebrate species to date^6^, and preliminary analyses by the authors of this study discovered some sequences and genes missing in previous reference assemblies of the same species^6^.

Here, in our companion study, we performed a more thorough evaluation of the VGP assemblies relative to previous popular references, when available. These species included a mammal (platypus), two birds (zebra finch, Anna’s hummingbird) and a fish (climbing perch). We found thousands of genes completely or partially missing as false gene losses in the previous assemblies, many located in newly identified chromosomes. The main causes for these false losses were the inability of short-read technologies to sequence through high GC-content regions, including the immediate upstream regulatory regions of the majority of genes with CpG islands, and difficulty in assembling repeat regions. These findings demonstrate the necessity of sequencing technology that sequences through GC-rich regions and generates long-reads longer than the repeat units in a genome, to obtain the complete gene landscape.

## Results

### Previously missing genomic regions have higher GC- and repeat-content

The VGP assemblies were on average ~635-fold more contiguous for contigs, in turn changing from ~200,000 to ~23,000 scaffolds in the previous assemblies to 100s in the VGP assemblies, for an expected 20 to 40 chromosomes (**Extended Data Table 1**). The VGP assemblies were also pseudo-haplotype phased, where paternal and maternal alleles were separated but assembled with switches between haplotypes. The zebra finch and Anna’s hummingbird previous and VGP assemblies were from the same individual animals, whereas the platypus and climbing perch were from different individuals. The prior zebra finch^12^ and platypus^13^ were chromosomal-level Sanger-based assemblies, whereas the Anna’s hummingbird^4^ and climbing perch^14^ were Illumina-based. To identify the specific sequence differences between the prior and the VGP assemblies, we aligned them using minimap2^15^ and cactus^16^; minimap2 provides better global alignments with higher stringency, whereas cactus is sensitive in detecting local synteny and can perform reference-free alignment between the prior assembly, the VGP primary assembly and the VGP alternate haplotype assembly. Both allow identification of sequences unique to each assembly. We took the intersection of findings between the two types of alignment as the most conservative estimate of differences between the prior and new assemblies (**Extended Data Table 1**). Genomic regions in the VGP assemblies with no alignment to the previous assembly were used as an approximation of missing regions, and vice versa. We removed the genes that were falsely duplicated due to haplotype separation errors or sequence errors in the VGP assemblies, that were identified in our companion study (Ko et al., submitted), so that they would not create false missing genes in the previous assemblies.

In all species, we found between 3.5 to 11.3% of genomic regions in the VGP assemblies were missing in the prior assemblies, affecting all chromosomes or scaffolds (> 100 kbp; **Fig. 1a-d**); this represented 37.5 to 213.4 Mb of missing sequence. However, the distribution was uneven, ranging from 1.2 to 96.7% per scaffold. We searched for a variable that could explain the cause of the missing sequences, and considered GC-content and repeat content, as suggested in Peona et al.^17^. We found that the higher the GC- or repeat content, the more missing sequence in the prior assemblies (**Fig. 1a-d**; **Extended Data Fig. 1a**). There were also many missing segments in the prior assemblies that had both higher GC-content and higher repeat content (**Fig. 1e**). However, the repeat content distribution was much wider than the GC-content distribution for the missing regions (**Fig. 1e**). Interestingly, the climbing perch had a more uniform and lower GC- and repeat content across all chromosomes, and likewise relatively uniform missing sequences in the previous assembly (**Fig. 1d**). Also, for the climbing perch, there was no difference in the GC-content of the previously missing sequence (**Fig. 1e**), consistent with a previous report of greater sequence homogeneity in fish genomes compared to tetrapods^18^.

**Fig. 1.**
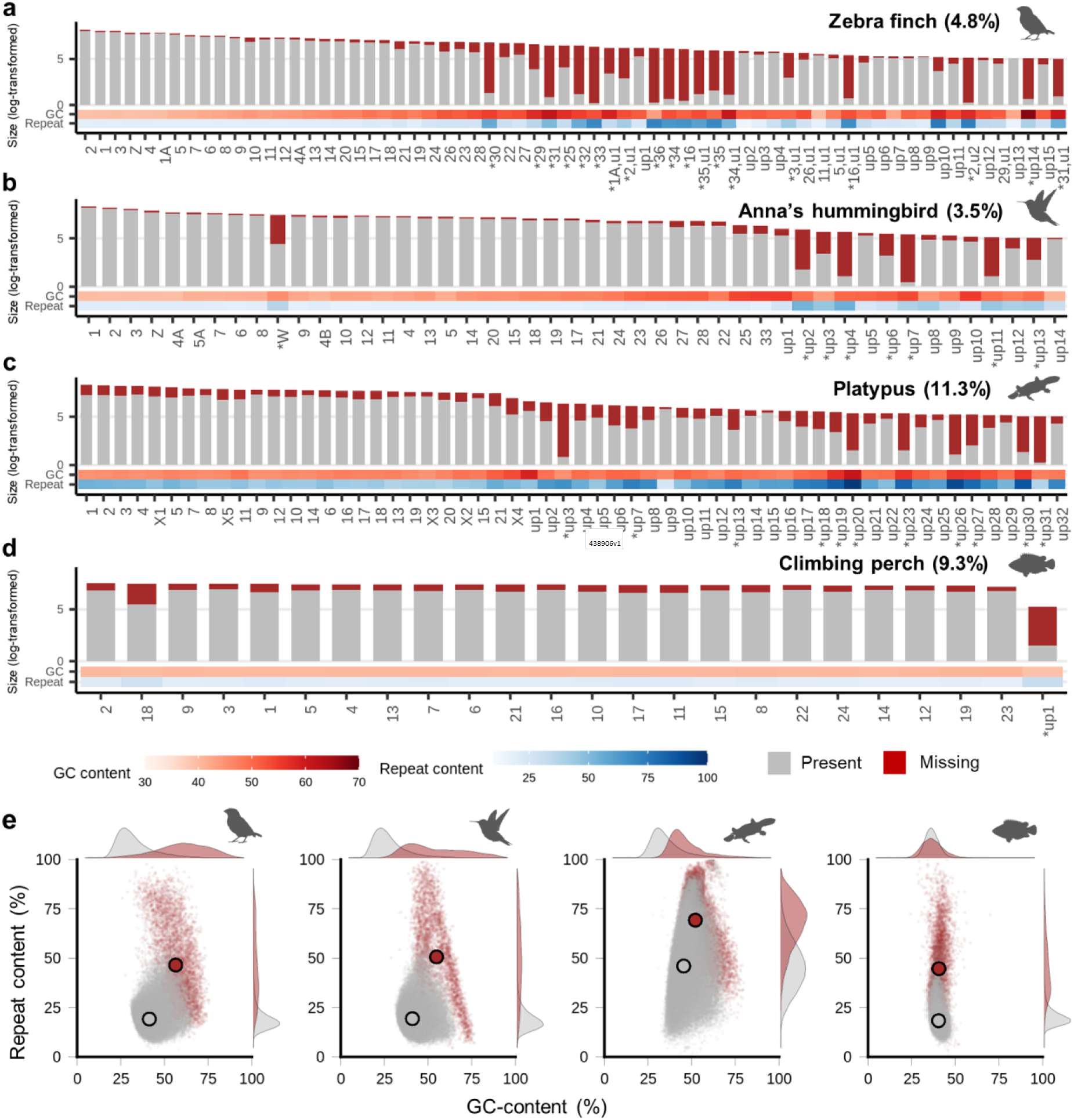
Proportion, GC-content, and repeat content of missing genomic regions in prior assemblies found in VGP assemblies. **a-d**, Logarithm of identified chromosome or scaffold size for those greater than 100 kbp in each of the VGP assemblies. Grey and red bars highlight the proportion of sequence present or missing in the prior assemblies, respectively. Below each chromosome/scaffold is a heatmap of the GC and repeat contents of the missing sequence. up, unplaced scaffolds; u, unlocalized within the chromosome named. * indicates the scaffolds with over 30% of missing sequences in the prior assembly. **e**, Distributions of % GC-content and % repeat content in 10 kbp consecutive blocks of missing or present sequences. Large dots indicate the average of GC and repeat content, which were significantly higher in the missing regions (red) than in the previously present (grey) regions except GC-content of climbing perch (p < 0.0001, Wilcoxon rank-sum test).

### Newly identified chromosomes are high in gene density

In the VGP zebra finch assembly, there were 19 scaffolds/chromosomes > 100 kbp that had over 30% (~36% to 97%) of their sequences entirely missing in the prior assembly (**Fig. 1a**, marked with *). Most of these scaffolds were originally considered as unplaced in our earlier curated version of the VGP assembly (GCA_003957565.2). We found that all were GC-rich, and had 400 genes completely missing in the prior assembly. This led us to reanalyze the Hi-C plots (**Extended Data Fig. 2**) using a more sensitive interaction map software than Juicer for smaller scaffolds, called PretextView (https://github.com/wtsi-hpag/PretextView) and HiGlass^19^. The maps revealed that 4 of the 19 scaffolds belonged to the ends of macrochromosomes 1A, 2, and 3, containing 19 of the completely missing genes (**Fig. 2a**, ‘ u’ for unlocalized scaffolds in thin purple bars); 3 of them greatly expanded the sizes of chromosomes 16 and 25 (by 1.33 Mb and 1.23 Mb, respectively), adding 47 of the completely missing genes (**Figs. 1a and 2b**); 11 belonged to 8 newly identified microchromosomes less than 6 Mb, containing 322 of the completely missing genes (**Fig. 2b**, purple bars. Chrs 29~36); and 1 still remains unplaced (up14). This newly curated assembly is GCA_003957565.3, updating the earlier VGP assembly. The 8 newly identified microchromosomes had similar GC and repeat content as the previously identified 4 microchromosomes (Chrs 16, 22, 25, and 27). The missing genes made up 0.1 to 1% of the genes in the macrochromosomes, but 6 to 50% of the genes in the microchromosomes. Consistent with these findings and a prior hypothesis^10,11^, the overall gene density was 3.7-fold higher in the microchromosomes < 10 Mb (40.8 genes per Mb) than in the macrochromosomes (10.8 genes per Mb; **Fig. 2** and **Extended Data Fig. 1b**). These findings indicate a preferential false loss of genes in the GC-rich microchromosomes of the previous zebra finch.

**Fig. 2.**
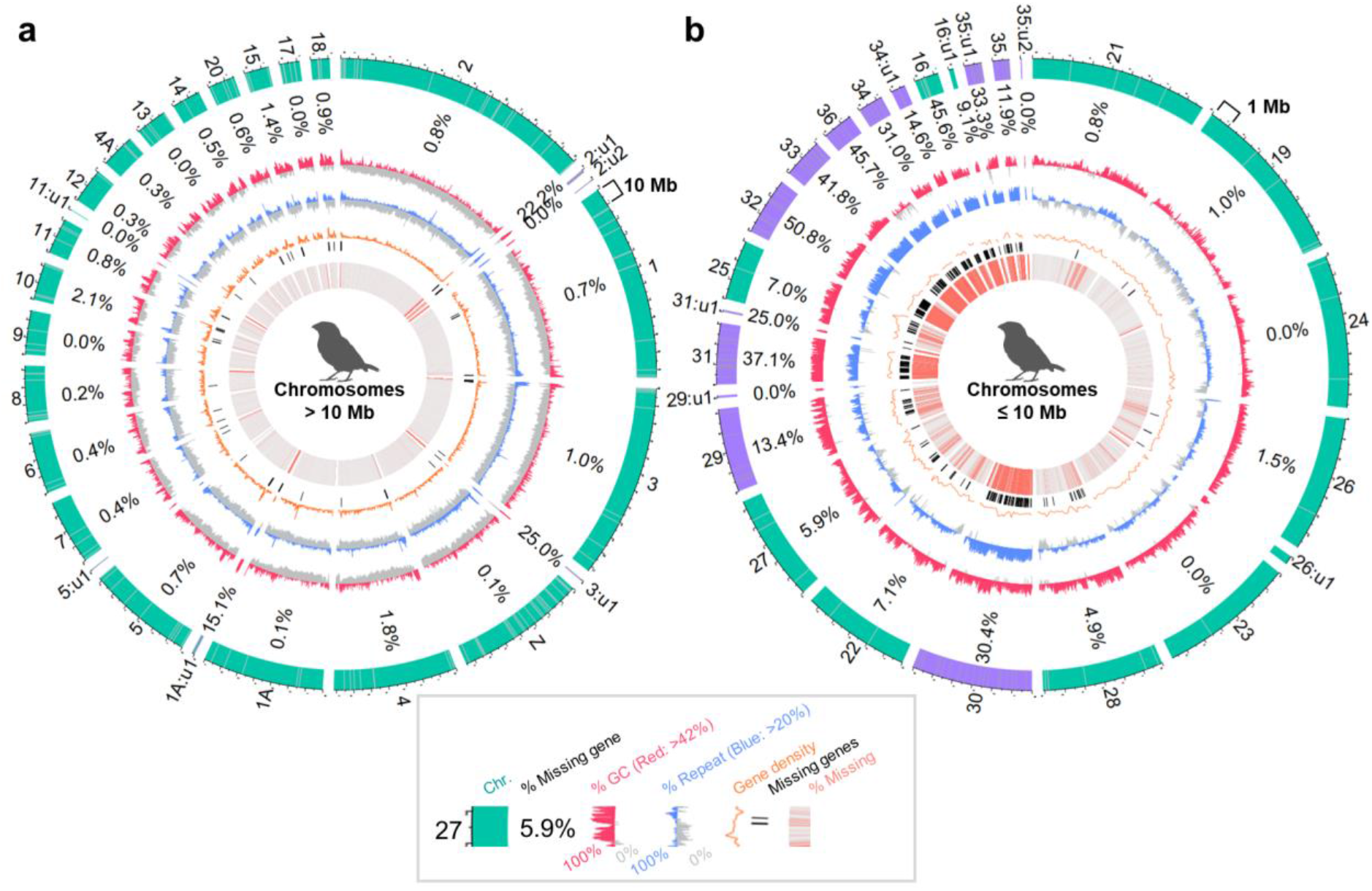
Circos plots revealing chromosome profiles of previously present and missing protein coding genes recovered in the VGP zebra finch assembly. **a**, Circos plot of chromosomes greater than 10 Mb in size. **b**, Circos plot of chromosomes less than 10 Mb in size. In the zebra finch, previously 20 Mb or 40 Mb were used to classify micro and macrochromosomes^22^, but we used 10 Mb for effective visualization. The two plots are not to scale. Shown from the outer to inner circle are: Chromosome number name (u: unlocalized) with previously present labelled in green, newly assembled and assigned labelled in purple, and assembly gaps labelled in grey lines in the outermost circle; % ratio of missing genes in the previous assembly; GC-content, over the average of 42% in red and under in grey; Repeat content, over the average of 20% in blue and under in grey; Gene density in non-overlapping 200 kbp windows, orange line; Loci of totally missing genes in the prior assembly, black bars; Alignment with the previous assembly, with red bars as unaligned regions. Circos plots were generated with R package OmicCircos^23^. Chromosome-level scaffolds were sorted in descending order by size. Each scaffold was binned in consecutive 10 kbp blocks. Missing ratio of protein coding genes was calculated by dividing the number of completely missing genes with the number of all genes on each scaffold. Gene density was calculated with BEDtools^24^ makewindows and intersect.

In the previous Anna’s hummingbird assembly, chromosomes were not assigned, but the amount of missing sequence we identified was less pronounced than in the zebra finch (**Fig. 1b**). This is presumably because it was generated with higher coverage (110x vs 6x). However, we still discovered that the smallest microchromosomes (Chrs 22, 25, 27, 28, and 33, all less than 6 Mb) showed higher missing ratios of 6.7-13.4% than the macrochromosomes (Chrs 1, 2, and 3, 1.7-1.8%), along with ~5% of higher GC content on average. Similar to the zebra finch, several GC-rich segments with high gene density were severely missing in the previous Anna’s hummingbird assembly (**Extended Data Fig. 3a,b**). Additionally, 40% of its W chromosome was missing in the previous assembly, which showed much higher GC- and repeat-content compared to similar-sized autosomes (**Fig. 1b**).

In the previous platypus assembly, chromosomes were assigned, but the VGP assembly newly assigned six chromosomes (Chrs 8, 9, 16, 19, 21, and X4) containing 2,834 genes in total. Although much of these sequences were previously assembled, they were too short in size to be scaffolded into chromosomes^6,20^. In the previous climbing perch assembly, chromosomes were not assigned, and sequence continuity was also too short. The VGP climbing perch genome assembly brought the sequences together into 23 chromosomes, consistent with the karyotype data^21^.

There are still 14-32 unplaced scaffolds > 100 kbp in the VGP zebra finch, hummingbird, and platypus assemblies, each containing sequences that were missing in the prior assemblies, half of which are GC-rich and repeat rich (**Fig. 1a-c**, up for unplaced). Like for the zebra finch, future higher resolution scaffolding data may identify additional smaller chromosomes or segments of currently identified chromosomes (**Extended Data Fig. 3c**), especially for birds. We believe that such smaller chromosomes could be difficult to identify with Hi-C short reads that do not sequence well through GC-rich regions or have mappability issues in highly repetitive regions. Small chromosomes are also difficult to identify in karyotyping. Our overall findings indicate that a large proportion of chromosomal segments or entire chromosomes were missing in prior commonly used reference genome assemblies, and these tended to be GC-rich, repeat-rich, and contain many genes.

### Bias of missing sequence in coding, lncRNA, and their regulatory regions

We asked if the relationship between the GC-content or repeat content and the missing sequence was randomly distributed among the genome or biased to protein coding genes. Supporting the later possibility, we found that the completely missing coding sequences in the prior assemblies were more GC-rich than those that were not missing, and only in the zebra finch they also included more repetitive regions (**Fig. 3a**). On average, 5 to 13% of the exons, introns, or intergenic regions had previously missing sequences, with birds having a higher proportion in the coding sequences (**Fig. 3b**). We compared the cumulative proportion of protein coding genes and discovered that ~83% of the genes had less than 10% missing sequence in the Illumina-based assemblies, whereas 56% and 77% of the genes had less than 10% missing sequence in the Sanger-based assemblies (**Fig. 3c**). That is, the genes in the Illumina-based assemblies were relatively well-assembled compared to the Sanger-based assemblies. In total, depending on species, between 3,479 to 20,132 exons in 14,648 to 23,833 genes were missing in the previous assemblies (**Supplementary Table 1**).

**Fig. 3.**
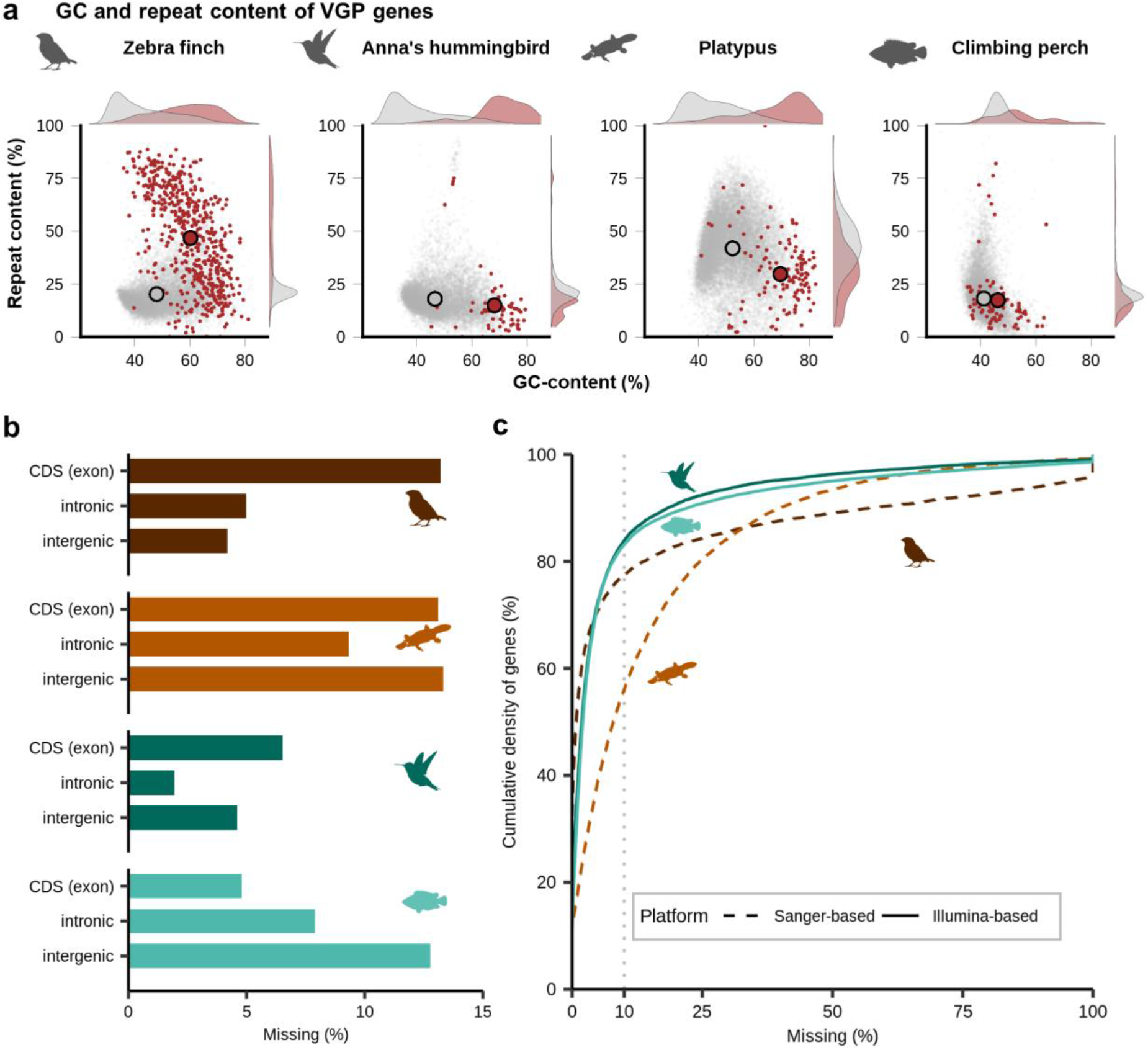
Amount and characteristics of missing genes and exons. **a**, GC and repeat content of completely missing genes in previous assemblies (red) compared to those of genes present (grey) within VGP assemblies. **b**, Percent missing of exonic, intronic, and intergenic sequences in the prior assemblies. **c**, Cumulative density plot of protein coding genes as a function of percent missing sequence. Illumina-based assemblies (Anna’s hummingbird and climbing perch) have more complete genes compared to Sanger-based assemblies (zebra finch and platypus). Grey dashed line indicates where 10% of a gene is missing.

Upstream of protein coding genes, we found an exponential increase in missing sequence in the birds and the platypus, changing from 10-20% missing at 3 kbp to 40-70% missing in the 100 bp window before the transcription start site (TSS; **Fig. 4a**). There was a similar higher missing percentage in the 5’ UTR and first exon, followed by a steady decrease in subsequent exons and the 3’ UTR until the transcription termination site (TTS). The percent missing sequence in the introns was much less than the exons, and the pattern was more stable between 5’ and 3’ introns. The fish were different in that there was not an increase in missing sequence closer to the TSS. The pattern of missing sequence within and across species was directly proportional to the GC-content (calculated in the VGP assemblies), in the promoter regions and for the UTRs, exons, and introns (**Fig. 4a**). This GC-pattern was biological, which we found in additional species sequenced in each vertebrate order in our companion study^6^. We further note here that the increased pattern of missing sequence was more prominent in the 75-80% of genes with upstream CpG islands in the birds and the platypus (**Fig. 4a**), further supporting a relationship between missing sequence and GC-content.

**Fig. 4.**
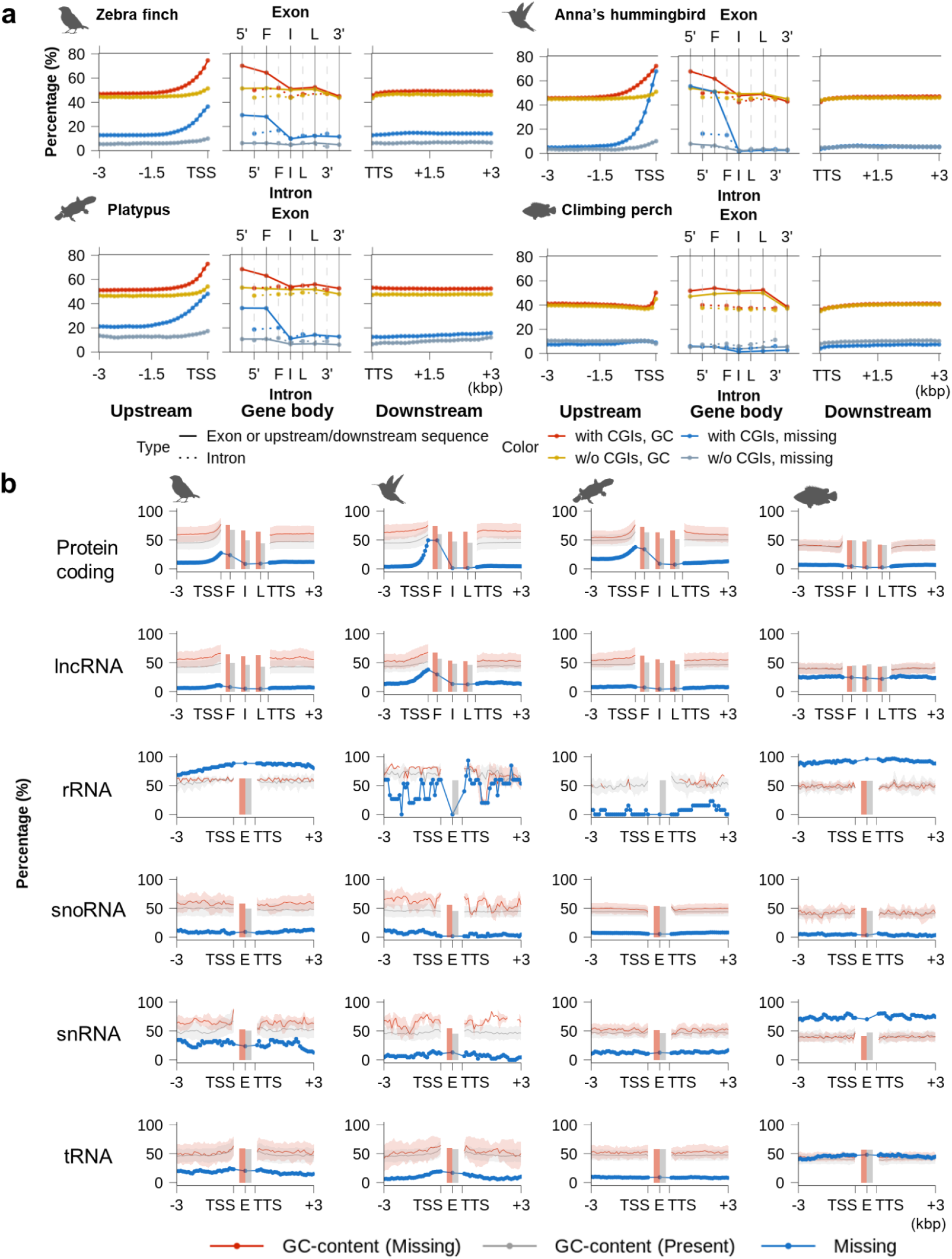
Distribution of missing sequences and GC-content within or near genes in previous assemblies relative to VGP assemblies. **a**, Average missing ratio and GC-content of VGP RefSeq annotated multi-exon protein coding genes separated by the presence or absence of upstream CpG islands (CGIs). Left and right panels indicate the upstream and downstream 3 kbp sequences of a gene in 100 bp consecutive blocks. Middle panels indicate the gene body regions with exons (top) and introns (bottom) positions. **b**, GC-profile of previously missing and present regions in various types of genes. Solid line with transparent background indicates average and S.D. of GC-content calculated from 100 bp consecutive blocks extracted from the upstream and downstream 3 kbp regions of genes. Blocks were classified as missing if their missing ratio was over 90%. Missing was calculated by the percentage of missing blocks among all blocks. Bar indicates the average GC-content of exons (F: first exon, I: internal exon, L: last exon, E: exon without consideration of its order).

We also found a similar pattern of missing sequence and GC-content for long non-coding RNA (lncRNA) genes and their regulatory regions (**Fig. 4b**). The other non-coding genes had 10-95% missing sequence, but without fluctuations in missing sequence or GC-content across the gene bodies (**Fig. 4b**). We believe their missing sequence is explained more by their repetitive nature (**Extended Data Fig. 4a,b**). We found an inverse relationship with repeat content and missing sequence surrounding protein coding genes, where the sequences were less repetitive around the beginning of the genes (**Extended Data Fig. 4c;** except climbing perch). These results demonstrate the dramatic impact of GC-content on missing sequences in coding and some non-coding genes, including their regulatory regions.

### False SNP and indel sequence errors in GC-rich regions

Single nucleotide polymorphisms (SNP) and insertions/deletions (indels) are some of the most valuable data for identifying sequence variation associated with specialized traits, genetic disorders, or phylogenetic relationships within and between species^25–27^. We asked if there were any false SNPs and indels due to sequence errors in the previous assemblies (those not explained by haplotype differences or sequencing errors in the VGP assembly), made possible to detect with the zebra finch and Anna’s hummingbird, since they were generated from the same individuals. We found false SNPs and indels, which like the missing sequences, were present in higher proportions in the 5’ proximal regions, and correlated with higher GC content (**Fig. 5a-c**). An exception was false indels in the prior Anna’s hummingbird assembly whose frequency did not increase in 5’ upstream regions but increased in UTRs and immediately downstream to the TTS (**Fig. 5d**). This difference appeared to be related to higher homopolymer content in the Anna’s hummingbird in the 5’ regions (**Extended Data Fig. 5a**), which is known as one of the main causes to induce indel problems in the Illumina platform^28^, suggesting a different error mechanism. These findings demonstrate that despite their levels of sequence accuracy in the prior Sanger- and Illumina-based assemblies, they have increased sequence errors around the beginning of protein-coding genes relative to the polished long read assemblies, and these errors lead to false SNPs and indels.

**Fig. 5.**
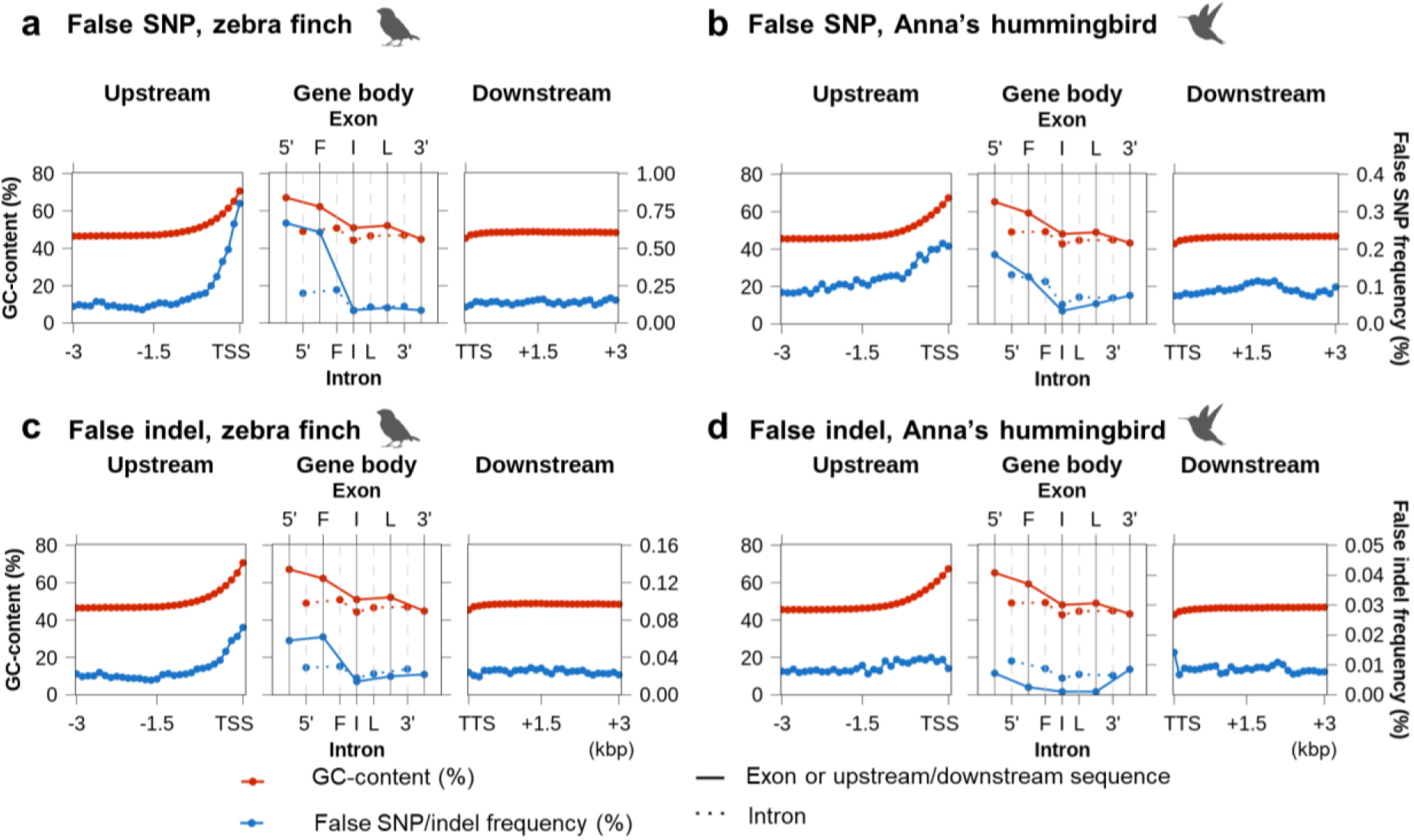
Biased distribution of sequencing errors near GC-rich 5’-proximal regions of protein coding genes. **a-d**, Average GC-content (red) and frequency of false SNPs or false indels (blue) found in the exons and introns of protein-coding genes (5’: 5’UTR, F: First coding, I: Internal coding, L: last coding, 3’: 3’UTR exon or intron). Left and right panels indicate the upstream and downstream 3 kbp sequences of genes in 100 bp consecutive blocks.

### Types of false gene losses in previous assemblies

Upon examining individual coding genes with false missing sequences, SNPs, and indels, and their annotations, we found that we could classify them into eight types of false gene losses: four types of structural errors (**Fig. 6a-d**) and four types of sequence errors (**Fig. 6e-h**). To quantify them, the annotation of the VGP assemblies were projected onto the previous assemblies using the Comparative Annotation Toolkit^29^ (CAT), and then these eight types of differences were searched for and quantified. We mapped Illumina reads from 10x Genomics libraries to both the VGP and previous assemblies and determined which assembly the reads had a mismatch, indel, or insufficient read depth for the sites nearby frameshifts, premature stop codons, and splicing junction disruptions (**Methods**).

**Fig. 6.**
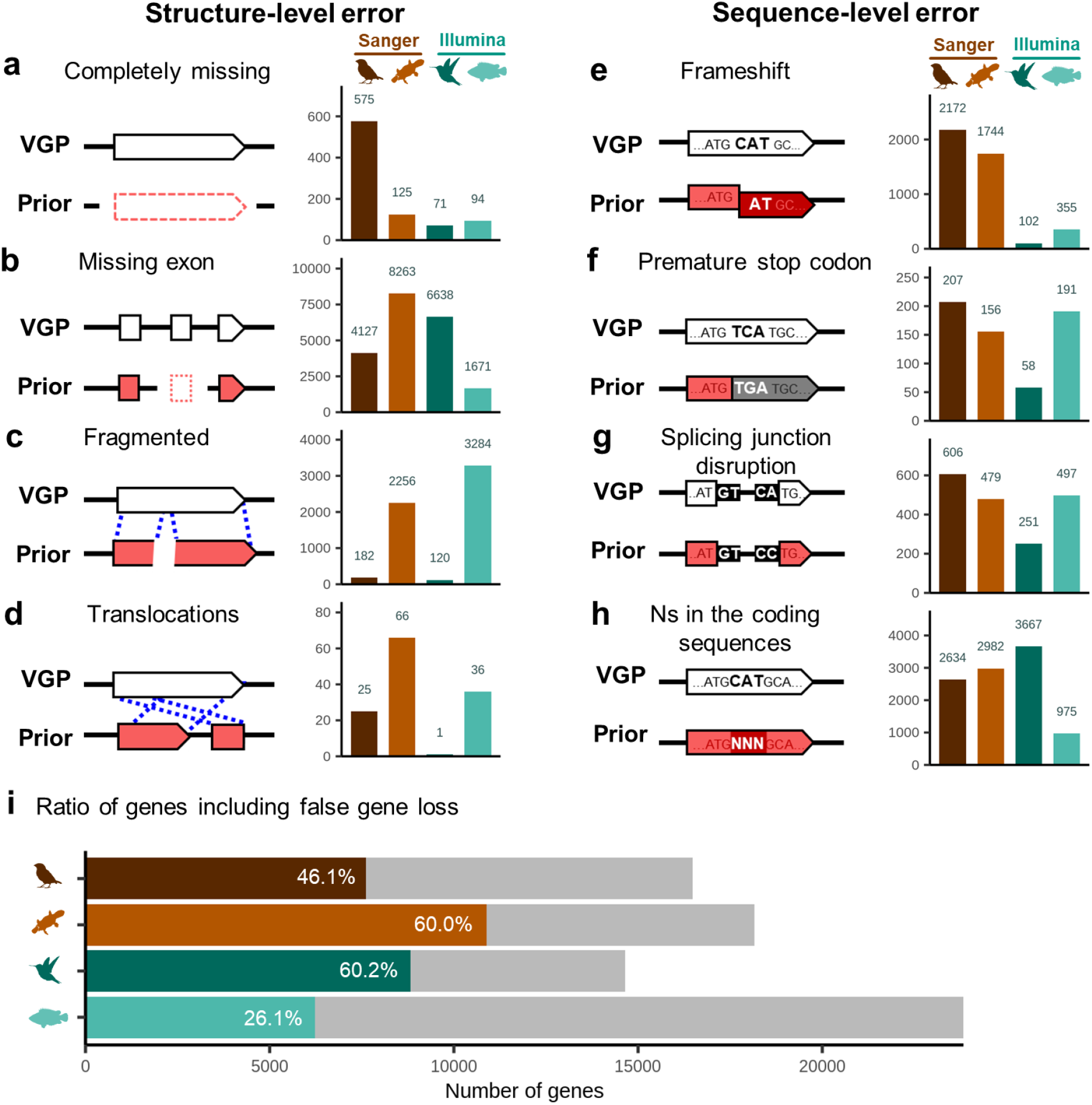
Types and amount of false gene losses in the previous assemblies relative to the VGP assemblies. **a-h**, Example model (left) and the number of genes affected in each species (right) by each type of false gene loss. **i**, Relative proportion (colored) of genes with false gene losses in the previous assemblies, calculated from the total number of annotated genes in the VGP assemblies (grey).

Depending on species, we found that a remarkable ~26 to 60% of genes in the previous assemblies contained one or more of these 8 types false gene losses (**Fig. 6i**). Missing exons (**Fig. 6b**) and Ns in the coding sequences (**Fig. 6h**) were the most frequently found false losses, while translocations (**Fig. 6d**) and premature stop codons (**Fig. 6f**) were the least. Even though the previous assembly of climbing perch included the least number of missing exons and Ns in the coding sequences, it included the most number (thousands) of fragmented genes (**Fig. 6c**), which is consistent with the fact that this assembly had the lowest contig NG50 (**Extended Data Table 1**). Consistent with lower frequency of indels in Illumina-based assemblies, the previous Anna’s hummingbird and climbing perch assemblies contained far fewer genes with frameshift errors (**Fig. 6e**). Overall, the substantial number of genes with missing or misrepresented sequence errors in previous assemblies highlight the importance of high-quality assemblies to provide more accurate gene information.

### False gene losses in previous annotation

Next, we tested the impact of GC-content on gene annotation. We compared RefSeq gene annotations on the previous and VGP assemblies (**Extended Data Table 1**). We also analyzed projected annotations by CAT from the VGP assemblies to the prior assemblies, to distinguish artifacts from version differences in the annotation process because the VGP assembly annotation was performed with updated gene models and more recent RNA-seq data. We note that RefSeq gene annotation was not available for the prior climbing perch assembly. Validating our analyses above, the GC-content of the annotated sequence rapidly increased before the TSS of the bird and mammal genes, and were overall ~2 to 15% higher in the VGP assemblies relative to the prior assemblies (**Fig. 7a**). We found a decrease in GC-content on the 3’ side of the TSS among all species (inclusive of exons and introns) forming an inverse pattern around the TSS site (**Fig. 7a**), consistent with previous observations^30^. Thus, the reduced GC-content function on the 3’ side of the TSS is smooth when measuring it in 100 kbp windows, but more step wise when analyzing UTRs and exons individually (**Extended Data Fig. 6a**). In contrast, we noted a smaller but present dip in GC-content on either side of the TTS (**Fig. 7a, right**). The GC-content of the projected annotations from the VGP to the previous assemblies were similar to the annotations of the previous assemblies (**Fig. 7a**), indicating that annotation version differences are not the main cause of the GC-content pattern differences. Instead, these results highlight the incomplete gene annotation in previous assemblies were due to limitations in sequencing GC-rich regions. On the other hand, the projected annotation of the prior climbing perch assembly showed similar GC content in the VGP annotation (**Fig. 7a, bottom**) since its genes have reduced GC-rich sequences relative to the platypus and the birds.

**Fig. 7.**
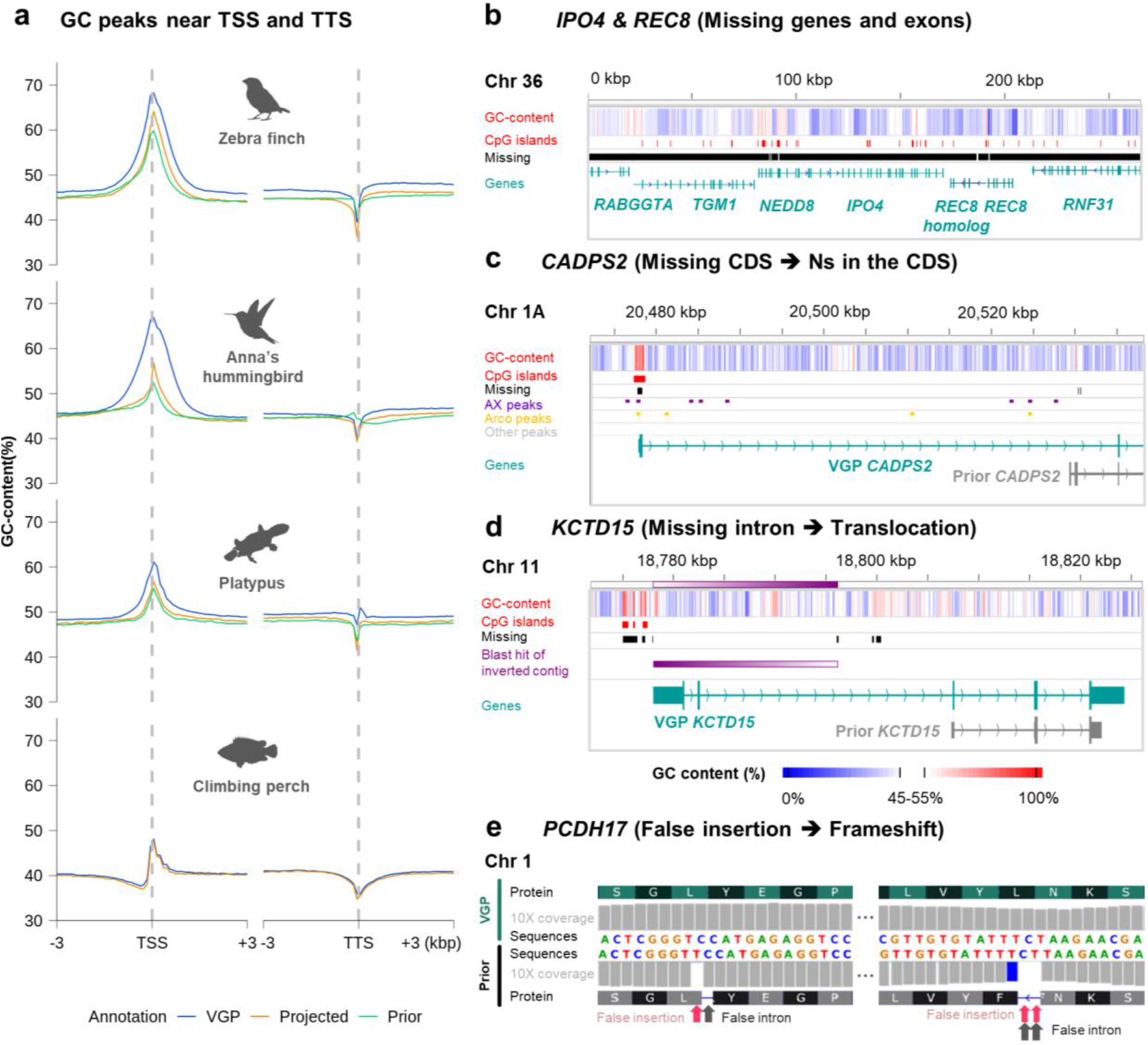
Effect of false gene losses in the previous assemblies on annotations. **a**, GC-content peaks near TSSs and TTSs from VGP or prior annotations (blue: VGP annotation, yellow: VGP annotation projected on the prior assembly by CAT, green: prior annotation). **b**, *IPO4* and *REC8* were present in the VGP zebra finch assembly while most of the exons were missing in the prior assembly. **c** *CADPS2* was missing its 5’ UTR and coding sequence, resulting in the false understanding of this gene’s structure previously and false annotation. **d**, *KCTD15* was erroneously assembled with the inverted contig including its first and second exons in the previous assembly. **e**, *PCDH17* included frameshift inducing indels in the coding region in the previous assembly, which resulted in false prediction of 1 and 2 bp length introns to compensate for the frameshift error.

To demonstrate how the missing sequences and incomplete annotations impact biological findings, we present a few examples found on the zebra finch, which had the most number of missing exons or genes in the prior assembly. The Importin-4 (*IPO4*) and meiotic recombination protein REC8 homolog (*REC8*) genes were previously reported as a candidate bird-deleted syntenic block^31,32^. However, we found these two genes on the newly discovered and assembled chromosome 36 of the VGP zebra finch assembly (**Fig. 7b**), consistent with findings in chicken that used long reads^33^. We found that only a single small 2,269 bp contig (NW_002223730.1) contained one of the twenty-nine exons of *IPO4* in the previous finch assembly. We also found *IPO4* was often partially assembled in other bird species assemblies that mainly used short reads from the Avian Phylogenomics study^4^, and included conserved synteny with nearby genes (**Extended Data Fig. 6b**). In the VGP assembly, all of the genes were greatly expanded in size. In another recent assembly using a VGP-like approach with long reads and long-range scaffolding on the Bengalese finch^34^, we also found these genes assembled and at larger sizes. This finding indicates that entire syntenic blocks of genes could be claimed as falsely missing in assemblies.

Calcium-dependent secretion activator 2 gene (*CADPS2*) regulates the exocytosis of vesicles filled with neurotransmitters and neuropeptides in neurons^35^ and shows specialized upregulated expression in several forebrain vocal learning nuclei of songbirds^36^. Thus, there has been interest in identifying the regulatory region responsible for this upregulation. We discovered a GC-rich 5’ exon and upstream regulatory region, the later with a differential ATAC-Seq signal in the robust nucleus of the arcopallium (RA) song nucleus versus surrounding neurons, that was missing in the prior assembly (**Fig. 7c**). This resulted in a false annotation of gene structure in the prior assembly, whereas the first non-GC-rich intron was misannotated as the regulatory region and two initial exons (**Fig. 7c**).

The potassium channel tetramerization domain containing 15 gene (*KCTD15*) is involved in formation of the neural crest^37^ and shows differential expression in song learning nuclei of songbirds^38^. We discovered that the GC-rich 5’ exons and introns were partially missing in the prior assembly, and the contig in between (ABQF01043589.1) was inverted and surrounded by gaps. Consequently, the first two exons on this contig could not be annotated (**Fig. 7d**). This finding indicates that gene structure can be problematic even if sequences are partially missing.

Protocadherin 17 (*PCDH17*) is also differentially expressed in songbird song nuclei^38^ and regulates presynaptic vesicle assembly in corticobasal ganglia circuits^39^. In the first coding exon of the previous *PCDH17* assembly of the zebra finch, we discovered false insertions in the GC-rich 5’-proximal exon (**Fig. 7e**). As a result, there were 1-2 bp false introns to compensate frameshift errors, leading to misrepresentation of the gene structure. This finding suggests that false indels can lead to misannotation of short, non-biological introns.

### Example of genomic regions falsely missing in a VGP assembly

We found that there were some sequences in previous assemblies not present in the VGP assembly. Some of these were false haplotype duplications in the previous assembly that were correctly prevented in the primary VGP assembly (see companion paper by Ko et al.). However, we discovered a few sequences in the previous assemblies that were incorrectly missing in the VGP assemblies (**Supplementary Table 2**). Some were due to read length cutoffs (> 10 kbp), where heavily GC-rich reads tended to be shorter or more repetitive and not incorporated into the initial contigs, as reported in our flagship paper^6^. To correct these, we lowered the read cutoff threshold in future assemblies. Another example we note here was a ~2.7 Mb region missing on chromosome 19 of the VGP zebra finch primary assembly (bTaeGut1_v1.p, GCA_003957565.2, **Supplementary Table 3**). When aligning the missing sequence found in the prior assembly to the VGP alternate haplotype assembly (bTaeGut1_v1.h, GCA_003957525.1), we found it was present as a false duplication in the alternate, across multiple contigs (**Fig. 8a**). Since the alternate assembly was not scaffolded into chromosomes nor annotated, it resulted in failure to annotate 46 genes in this region in the primary assembly. The missing region was present as 2.7 Mb in the VGP assembly of another female zebra finch paternal haplotype (bTaeGut2.pat.W.v2, GCA_008822105.2), whose haplotypes were separated with trio-binning^40,41^ using parental data. We estimated repeat content in consecutive 10 kbp blocks extracted from the missing 2.7 Mb region and whole genome of bTaeGut2.pat.W.v2, and found that this missing region was significantly more enriched with LINEs and LTRs compared to repeat content of the genome of bTaeGut2.pat.W.v2 (**Fig. 8b,c**, ANOVA, p < 0.0001). These and other findings of some remaining false gene duplications (Ko et al., submitted) led us to reassemble the VGP male zebra finch assembly using improved algorithms that we generated since the initial assembly (GCA_003957565.4 pending). However, the missing sequence was again found only in the alternate assembly. This highlights that even though the VGP assemblies recovered more GC and repeat rich sequences, still some of the genomic regions highly enriched with repeats are still a challenge and needs to be improved.

**Fig. 8.**
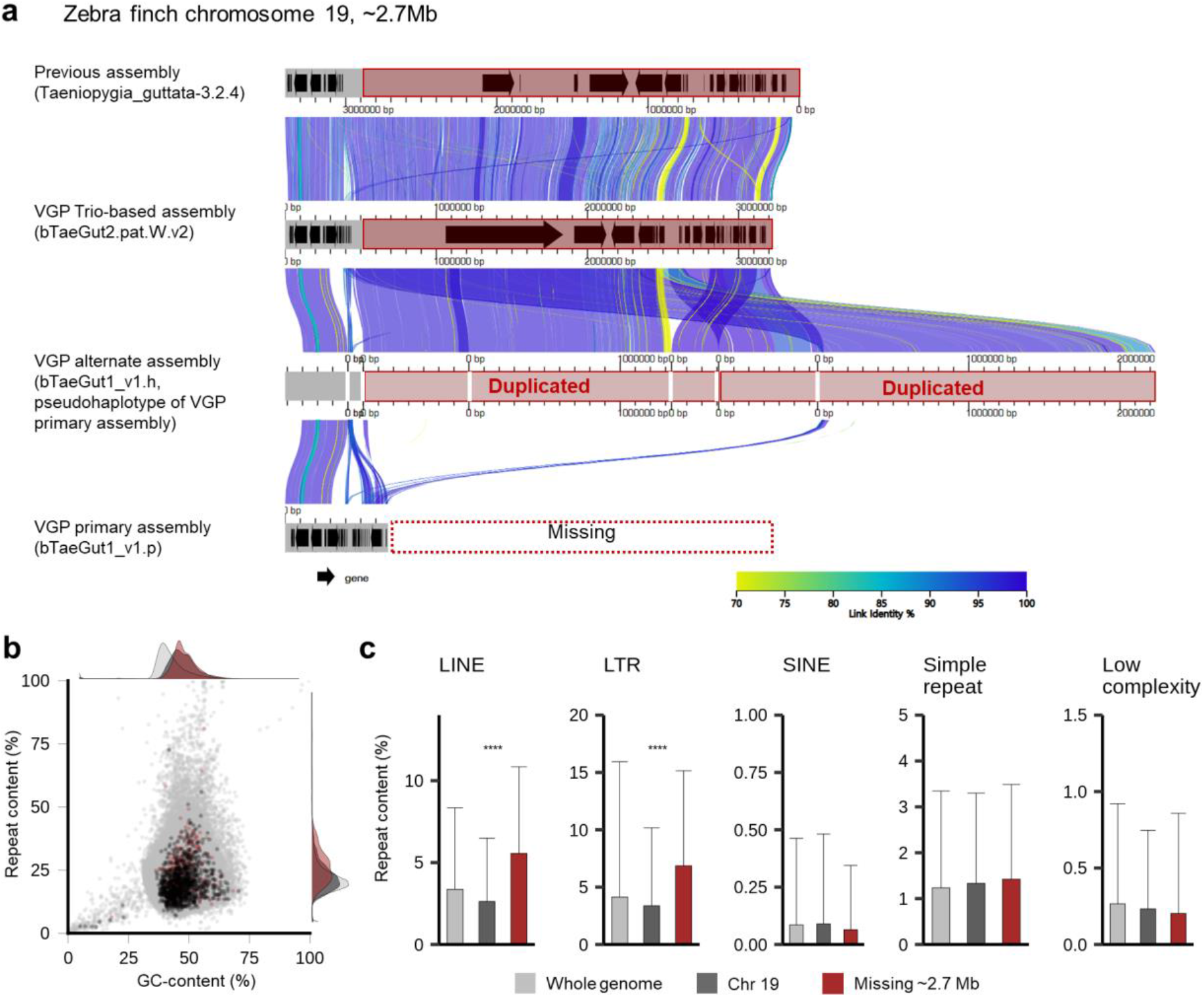
Genomic regions that failed to be assembled in chromosome-level scaffolds of the VGP zebra finch primary assembly (bTaeGut1_v1.p). **a**, Alignment between the previous, VGP Trio-based, VGP alternate and VGP primary assemblies for a 2.7 Mb end of chromosome 19. Grey, chromosome-level scaffolds. Black arrows, annotated genes. Links between grey bars indicate the alignment between each scaffold. **b**, GC- and repeat content of the 2.7 Mb region missing in the VGP primary assembly. Grey, dark grey, and red indicate GC- and repeat content calculated from 10 kbp consecutive blocks extracted from the whole genome of a VGP trio-based assembly, chromosome 19, and the 2.7 Mb end of chromosome 19, respectively. **c**, Repeat profile of the 2.7 Mb region missing in the VGP primary assembly. Repeat content was calculated from 10 kbp consecutive blocks extracted from the whole genome (grey), chromosome 19 (dark grey), or 2.7 Mb end of chromosome 19 (red) of the VGP Trio-based assembly. Bars and error bars indicate the mean and S.D. of repeat content of the blocks (****: p < 0.0001, ***: p < 0.001, **: p < 0.01, *: p < 0.05. P-values were calculated by ANOVA).

## Discussion

We find that previous reference assemblies based on short or intermediate read lengths miss or misrepresent many genes and even chromosomes. Besides short read lengths making it difficult to assemble repetitive regions, another major issue is difficulty in sequencing through and representing GC-rich regions. This led to false gene losses, false SNPs, or false indels, surprisingly biased to proximal regulatory regions of coding genes and lncRNA genes, and their 5’ exons and UTRs in more than half of the genes in some species. We did not know that such regions existed without having the new VGP assemblies. We also discovered completely missing genes, many of which were considered lost or “hidden genes” not assembled in birds^42,43^. These findings also led us to reveal a canonical pattern of GC-content in protein coding and lncRNA genes, advancing previous studies^44–46^.

Peona et al.^1^ reviewed the status of the completeness of prior bird genome assemblies generated with intermediate and short reads, by comparing estimated genome size and the size of the genome assemblies. Their estimates of the amount of missing DNA was 0.9% and 23.6% in the previous zebra finch and Anna’s hummingbird assemblies, respectively. However, compared to the VGP assemblies, we found that the amount missing was 4.8% and 3.5%, respectively. For the zebra finch, NCBI’s remap alignment between the VGP and previous zebra finch assembly showed 5.3% (56.3 Mb) of VGP sequences without any hit against the previous assembly, which contains 84% of the missing sequence we identified. For Anna’s hummingbird, we think Peona et al.’s calculations were in error due to miscalculations in genome size as 1.41 Gbp. Based on animal genome size database^47^ and its reference^48^, we estimate the genome size to be 1.11 Gbp (C value conversion into Gbp where 1 pg = 0.978 Gbp^49^), consistent with our other genome-size estimate of 1.12 Gbp using a k-mer based calculation of the raw sequence reads^6^. If we use this newly estimated genome size for Peona et al.^1^’s missing ratio calculation, almost no region is expected to be missing. We think the reason for our higher estimate of missing sequence has to do with false duplications we found in the 3.7-15.9% of the previous assemblies^2,6^ (Ko et al., submitted). False duplications are more prevalent when the assemblies are not haplotype phased (previous zebra finch assembly) or partially phased (previous Anna’s hummingbird assembly). Additionally, in our companion paper^6^, we found that substantial amounts of genomic regions were missing in both prior assemblies based on k-mer completeness estimation. Thus, we believe our estimates of the amount of missing sequences in the prior assemblies are more accurate.

Our finding of higher GC-content in the missing regions of the prior assemblies is consistent with GC-rich regions having been a challenge for both Sanger and Illumina platforms^50,51^ due to requirements of higher melting temperature in the sequencing process involving PCR or formation of a secondary structure^52^. This leads to skewed coverage in GC-rich regions (GC-bias^53^) and increases the probability of missing or misrepresentation of these regions. The presence of these GC-rich sequences in the VGP assemblies is related to the successful ability of the PacBio platform to sequence GC-rich regions^54,55^. What we find striking is that so much of the so-called “dark matter” missing in prior genome assemblies^56^ are parts of protein coding genes and their immediate upstream regulatory regions. Such recovery makes it difficult to justify generating whole genome assemblies with approaches that do not get through these GC-rich regions.

Our finding of higher repeat content in the previously missing regions is expected and consistent with the ability of longer reads and long-range scaffolding to sequence through and resolve them in the VGP assemblies^2,6^. Although this resulted in greater resolution of repeats, the results here and in our companion study^6^ suggest the possibility that species-specific repeat profiles may actually remain difficult to assemble into chromosome-level scaffolds even in the VGP assemblies. Ongoing improvements to long-read sequencing technologies and algorithms are being developed, such as for trio-binning^40,41^, PacBio’s circular consensus sequencing platform^57^ (CCS or HiFi), and Nanopore’s v10.3 chemistry^58^.

In conclusion, we identified and classified a significant amount of false loss present in the previous genomes of vertebrates. These false losses impacted many genes, regulatory regions, and their annotations. Fortunately, it was possible to recover these false losses in the VGP assemblies, which were done with long reads that can read through repetitive and GC-rich regions, long-range scaffolding data, and new algorithms that include haplotype phasing.

## Methods

### Genome alignment and analysis on the missing genomic regions

We analyzed zebra finch, platypus, Anna’s hummingbird, and climbing perch genome assemblies that were available in two versions: VGP 1^st^ release and a previous assembly generated from Illumina or Sanger sequencing platforms. The assemblies were downloaded from NCBI RefSeq^59^ and GenBank^60^, including the alternate haplotype for the VGP assemblies (**Extended Data Table 1**). We removed the mitochondrial genome associated with the assembly in order to prevent misalignment between mitochondrial and nuclear genomes. We then aligned the previous, VGP primary, and VGP alternate assemblies with cactus^16^, a reference-free genome-wide alignment tool. In order to define the regions that were not aligned by cactus, we used halLiftover^61^ (v2.1) to obtain the coordinates of the aligned regions in VGP assemblies and BEDtools^24^ (v2.27) subtract to exclude these regions. Additionally, we performed a minimap2^15^ (2.17-r974-dirty version) alignment with the following command: *minimap2 -x asm5 -r 50 -c -g 1000 -t 30 --no-long-join [VGP assembly] [Prior assembly]*. With paftools (bundled with minimap2) and BEDtools, we obtained the unaligned regions by minimap2. The coordinates of missing regions of each assembly were defined by the intersection of unaligned regions of both cactus and minimap2 alignment results (**Extended Data Table 1**). We took this conservative approach, because in the generation of the initial graph from cactus alignment, softmasked regions are excluded and only the unmasked regions are used for the self-alignment and pairwise alignment. Even though the masked regions between the unmasked regions can be recovered by the extension of the alignment from anchors, it can result in lower alignment ratio in the highly softmasked regions. The minimap2 alignment uses a different approach, by avoiding high frequency minimizers in seeding, alleviating a biased alignment. Alignments to the VGP assembly Y chromosome were excluded in the platypus, since the previous assembly was generated from a female while VGP assembly was from a male. Genomic sequences from the VGP assemblies identified as missing or present in the prior assembly were extracted and concatenated to calculate GC and repeat content in each 10 kbp non-overlapping window.

### GC-content and repeat content

GC-content was calculated by the summed number of Gs or Cs divided by the size of given coordinates excluding ambiguous nucleotides (N) with BEDtools nuc. Repeat content was calculated by extracting the coordinates of softmasked regions by WindowMasker^62^ and using BEDtools intersect in order to count the number of softmasked nucleotides.

### Analysis on the missing genic and exon regions

To estimate the relative amount of missing sequences in intergenic, intronic, and exonic coding sequences, for each VGP assembly, we downloaded GFF annotation file from NCBI RefSeq database and chose the longest transcript as representative of each gene (**Extended Data Table 1**). The VGP genes that were found to be falsely duplicated (Ko et al., submitted) were excluded. Using gffutils (https://github.com/daler/gffutils), coordinates of each region were extracted from the filtered GFF file and merged with BEDtools merge for the VGP assemblies. The length of intersection between the coordinates and the previously missing regions was calculated by BEDtools intersect and divided by the summed size of the coordinates to calculate the relative ratio of previously missing sequences. The function stat_ecdf from R package ggplot2^63^ was used to generate the cumulative density plot based on the missing ratios of VGP genes.

Given that some genes or exons present in the previous assemblies may not align due to limitations in the genome-wide alignment tools, we also performed blastn 2.6.0+ alignment^64^ of the VGP coding exons over 15 bp against the previous assembly with the following options: *-task blastn, -perc_identity 90, -qcov_hsp_perc 50, -dust no*, and *qcovus*. Blast hits with *qcovus* over 90% were considered reliable, which is a measure of query coverage which counts a position in a subject sequence for this measure only once^65^. A gene or exon in the VGP assembly was classified as completely missing when it had no cactus or minimap2 alignment and no blast hit against the previous assembly for its exons.

### Analysis of missing sequences in coding and non-coding genes

We regarded upstream and downstream 3 kbp regions to include potential regulatory regions of all genes^66^. To classify protein-coding genes based on the presence of CGIs, EMBOSS (version 6.6.0.0) newcpgreport^67^ was used to detect the CpG islands in the VGP assemblies with default settings: at least 200 bp length, a GC-content greater than 50%, and observed-to-expected CpG ratio greater than 0.6. The distance of the nearest upstream CpG island to each gene was calculated by using BEDTools closest with the following options: -id for ignoring 3’ downstream distance and –D for the bed file of the most 5’ proximal exons. Protein coding genes with four or more coding exons were classified into five categories: 5’ UTR, 3’ UTR, first coding exon, internal coding exon(s), and last coding exon. Exons annotated as both UTR and coding sequences were divided into two separated coordinates. The coordinates of 5’ UTRs, 3’ UTRs, coding exons and introns, and consecutive 100 bp blocks from the 3 kbp upstream of the TSS and 3 kbp downstream of the TTS were extracted by a custom Python code (https://github.com/JuwanKim67/FalseGeneLoss). For non-coding genes, we used exons from lncRNA genes, rRNA genes, snRNA genes, snoRNA genes, and tRNA genes. The blocks and exons were classified into missing if 90% or more of the region was missing in the alignment. Missing ratio of each block in the alignment was calculated by dividing the number of genes with missing blocks by the number of all genes at each position. Average and standard deviation of GC-content of each block was calculated.

### False SNP and indel variants

To call the false SNPs or indels from the previous assembly (**Extended Data Fig. 7**), we transformed the hal file from the cactus alignment to a variation graph using the hal2vg (v2.1) and vg toolkit^68^ (v1.27.1) with the following command: *hal2vg --inMemory --progress --noAncestors*. We called all variants using the VGP primary assembly as a reference using the following commands: 1) *vg index [*pg file*] -x [*xg file]; and 2) *vg deconstruct -e -v -P [VGP primary assembly] -A [Prior assembly] -A [VGP alternate assembly]*. In order to split multi-nucleotide variants into several single nucleotide variants, custom Python code was used for the normalized VCF file generated by the following command: *bgzip -c & bcftools index & bcftools norm -m –any.* The variant specifically found in the previous assembly was selected by the following command: *bcftools view -i’ (COUNT(GT[*Prior assembly*] = “alt”)> 0 & COUNT(GT[*VGP alternate assembly*] = “alt”) = 0)’ -v [snps* for SNP*, indels* for indel]. We excluded variants with more than one alignment within the prior assemblies. Using a custom Python code, the candidates of false SNP and indel variants under 10 bp between the VGP primary and prior assemblies were selected. The VCF files with the candidate false SNPs and indels were converted to BED files by vcf2bed. In order to exclude variants with VGP assembly base call errors or alternate haplotypes, we performed SAMtools (version 1.9-177-g796cf22) mpileup^69^ with the following command: *samtools mpileup -Bx -s - aa --min-BQ 0* and listed the loci with more than 10 reads and more than 20% of reads supporting the insertion, deletion, or mismatch (**Extended Data Fig. 7,8**). The candidate false SNPs and indels with any intersection with the flanking 2 bp of the loci of potential VGP erroneous sites or heterozygous sites were excluded. Homopolymers were defined as five or more consecutive stretches of the same nucleotide. As in the missing ratio analysis, the frequency of false variants was calculated by dividing the total number of genes with blocks containing the false variants by the number of all genes at the same position.

### Classification and detection of false gene loss candidates

We used the Comparative Annotation Toolkit^29^ (CAT), a program that projects annotations from a reference assembly to a target assembly based on a genome-wide alignment performed by cactus. We utilized CAT to project the VGP annotation to the previous assembly. Using a custom Python code, we classified false gene loss into eight types based on the following criterion:

1. **Completely missing:** Genes in one assembly with no genome-wide alignment nor exon-wide blast hit in the other assembly.
2. **Missing exon:** Genes in one assembly without any alignment of one or more exons and no exon-wide blast hits in the other assembly.
3. **Fragmented:** Genes labelled as “possible_split_gene_locations” in the CAT annotation, and where one or more exon blast hits had overlap in the coordinates of the projected gene and its split locations on different scaffolds in the prior assembly.
4. **Translocation:** Genes labelled as “possible_split_gene_locations” in the CAT annotation alignment and where one or more exon blast hits overlap with both the coordinates of the projected gene and its split locations that are on the same scaffold.
5. **Frameshift:** We performed blat^70^ (v. 36×2) using a database of the annotated VGP coding sequence and a projected coding sequence as a query for each gene, to detect the position of indels. Based on the output PSL file, we regarded the case as a frameshift if there was an indel whose length was not multiples of three and if the size of the indel was less than 10 bp for either the VGP or previous assembly.
6. **Premature stop codon:** Centered on the previous assembly’s coordinates of the “InframeStop” label, five bases of flanking sequences were taken and halLiftover was performed. To check whether the region in the VGP assembly was the same genomic region of the previous assembly, the alignment blocks within the same gene on both assemblies were first selected; only the case with identical flanking sequence was regarded as a candidate for a premature stop codon.
7. **Ns in the coding sequences:** Cases in which transcripts annotated by CAT included Ns in the coding sequences in the prior assembly.
8. **Splicing junction disruption**: The VGP assemblies’ splicing donor and acceptor sequences with flanking five bases were lifted to the previous assembly by halLiftover. We regarded introns of the VGP and previous assembly as the same, only when there was a single intron between the lifted donor and acceptor sites in the previous assembly. Canonical splicing junction sequences (one of GT-AG, GC-AG, or AT-AC) in the VGP and previous annotated introns were compared and classified as a splicing junction disruption in cases where the VGP assembled gene was misrepresented in the previous assembly. Cases with the prior splicing junction sequences with ambiguous nucleotides (32.5~82.9% of the introns with non-canonical splicing junctions, **Supplementary Table 4**) were excluded.

The candidate erroneous sequences of frameshifts, premature stop codons, and splicing junction disruptions were further filtered by mapping 10X genomics reads collected from the GenomeArk in github (https://vgp.github.io/genomeark/). 10x barcoded read mapping to both the VGP primary and the prior assemblies were performed with EMA^71^ (v0.6.2, https://github.com/arshajii/ema), following the pipeline in github page as below:

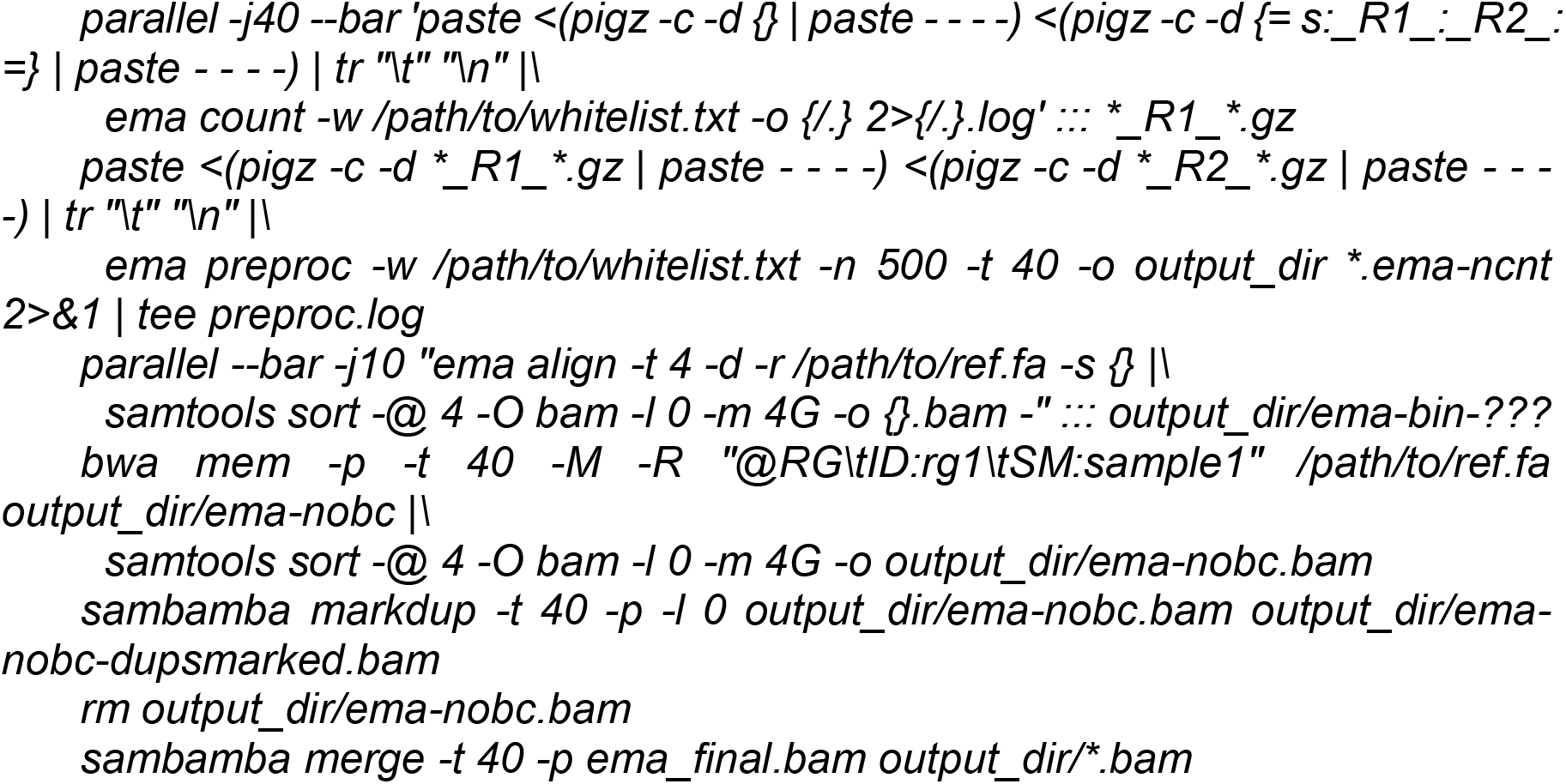

After read mapping, flanking two bases of each sequence-level false gene loss were parsed from the BAM file. SAMtools’ mpileup was performed to sum up the number of mapped reads on each locus with the following options: *-Bx -s -aa --min-BQ 0*. For premature stop codon and splicing junction disruptions, loci with fewer than ten reads aligned or with 80% of aligned reads containing mismatch or indels were regarded as assembly errors. For frameshifts, loci with the 80% or more reads containing indels were regarded as assembly errors (**Extended Data Fig. 9a,b**).

### Discovery of missing regions in the VGP zebra finch assembly

We performed halLiftover from the previous to the VGP zebra finch assembly and minimap2 alignment with the following command: *minimap2 -x asm5 -r 50 -c -g 1000 --no-long-join [Prior assembly] [VGP assembly]* and took the intersection of unaligned regions by each aligner. To get the sequences specifically found in the prior assembly, we excluded false duplication in the prior assembly from the unaligned regions (Ko et al., submitted). We took the longest missing sequence, the end of chromosome 19, to check whether this region was truly missing in the VGP zebra finch assembly (bTaeGut1_v1.p). We performed genome-wide alignments between the VGP primary (bTaeGut1_v1.p) or alternate haplotype (bTaeGut1_v1.h) assembly and the trio-based bTaeGut2.pat.W.v2 assembly, with both cactus and minimap2. In order to visualize the alignment between the end of chromosome 19 in bTaeGut1_v1.p, bTaeGut1_v1.h, the previous assembly, and the bTaeGut2.pat.W.v2 assembly, the fasta sequences of the end of chromosome 19 in each assemblies were extracted (**Supplementary Table 3**). The sequences were aligned and visualized with AliTV^72^. In order to investigate repeat content, consecutive 10 kbp blocks were extracted from the missing ~2.7 Mb sequence, whole chromosome 19 sequences, and whole genomic sequences of bTaeGut2.pat.W.v2. Their repeat content was then calculated with RepeatMasker^73^ output of bTaeGut2.pat.W.v2 from NCBI RefSeq by BEDtools intersect and groupby. ANOVA test was performed by R package ggpubr (https://github.com/kassambara/ggpubr) to compare repeat content of the ~2.7 Mb end of chromosome 19, that of entire chromosome 19, and that of the entire sequence of bTaeGut2.pat.W.v2.

## Extended Data

**Extended Data Table 1.**
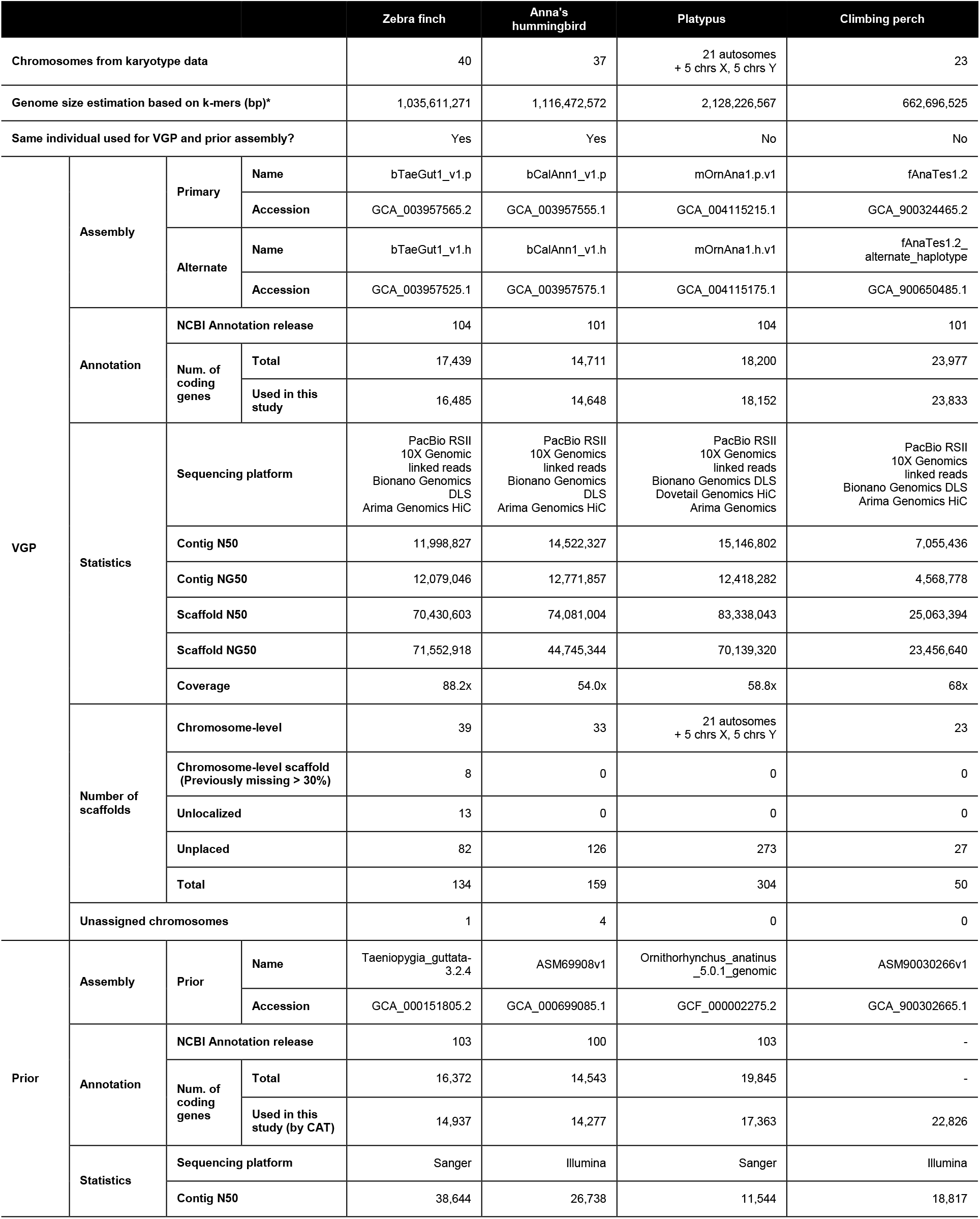
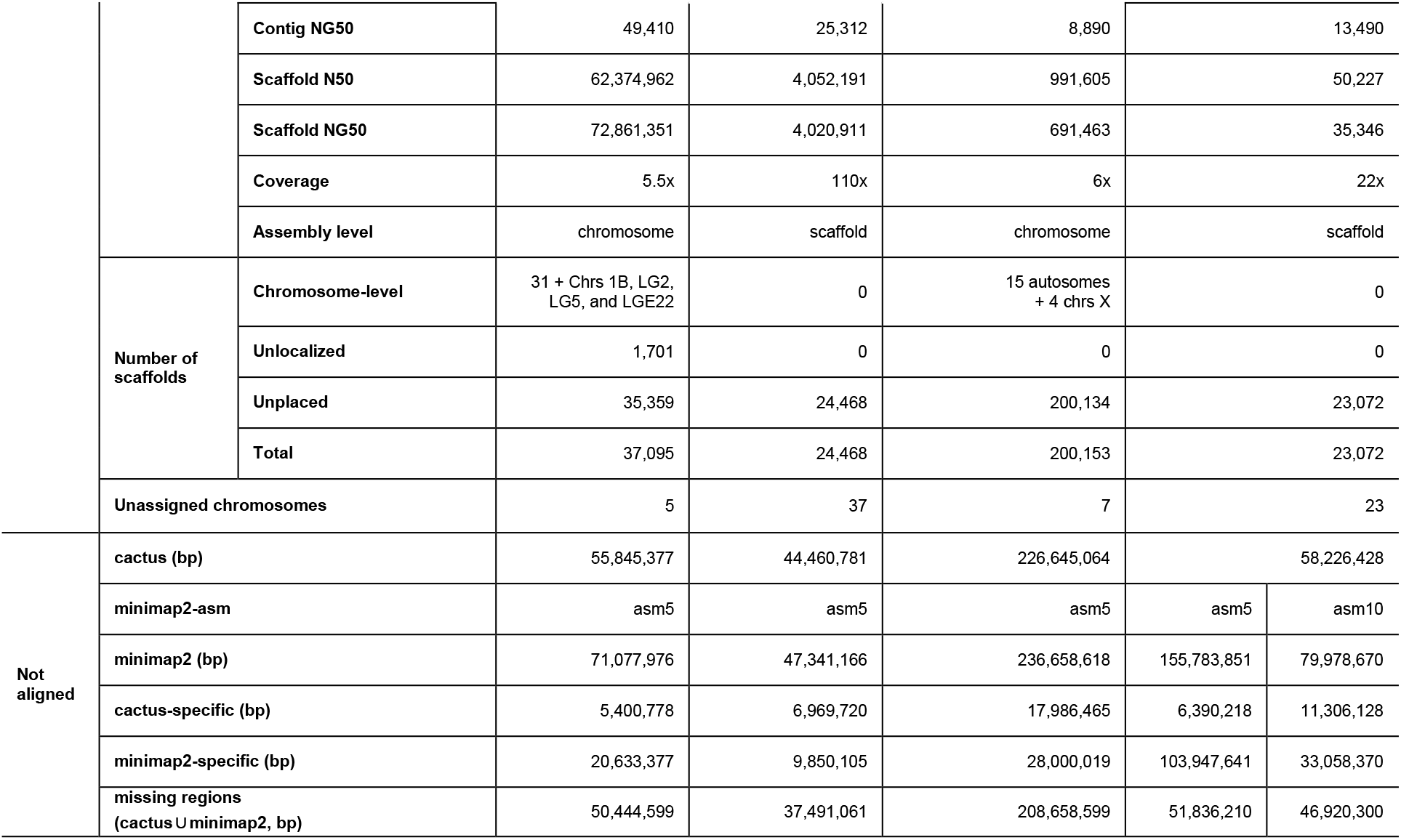
Assembly and annotation information used in this study. VGP: assembly and annotation information of the VGP assemblies. Prior: assembly and annotation information of the prior assemblies. Not aligned: the length of unaligned sequences from each aligner calculated based on the VGP assemblies. For minimap2 alignment of the climbing perch, two different options (asm5 for 5% of maximum sequence divergence, asm10 for 10% of maximum sequence divergence) were tested since the minimap2 result showed much larger estimation of unaligned regions compared to cactus. *: Genome size estimation was from Supplementary table 11 from the VGP flagship paper^6^.

**Extended Data Fig. 1.**
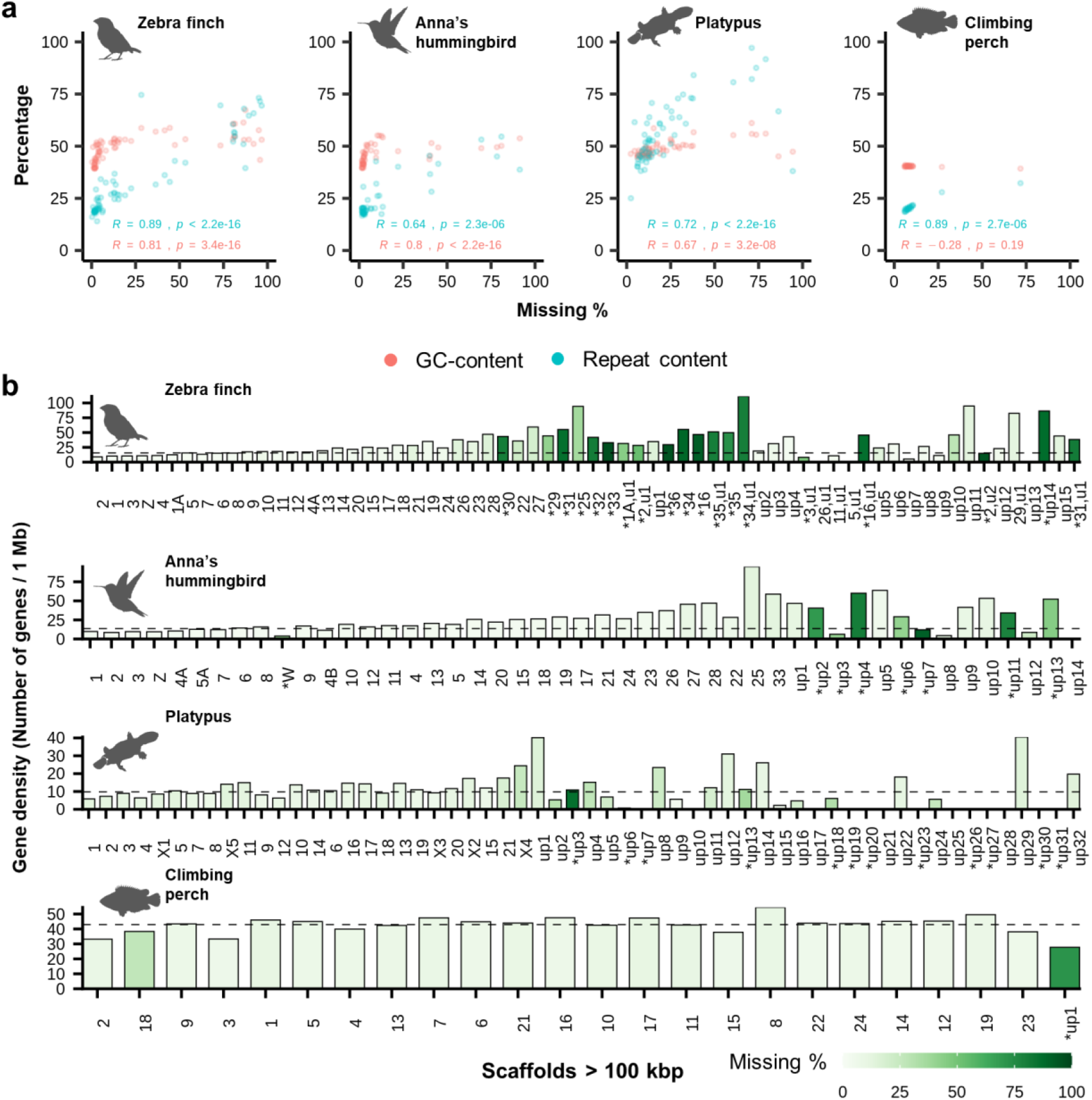
Association between missing ratio, GC- or repeat content, and gene density of VGP assemblies. **a**, Correlation between missing ratio and GC- or repeat content of VGP scaffolds > 100 kbp. Each dot represents the value of GC (red) or repeat (blue) content of each VGP scaffold and its missing ratio in the previous assemblies. Spearman correlation coefficients were calculated by R. **b**, Missing ratio and gene density of VGP assembled chromosomes. Bars indicate gene density of VGP chromosomes. Black dashed line indicates the average gene density. Chromosomes/scaffolds are ordered from largest to smallest, as in Fig. 1 a-c.

**Extended Data Fig. 2.**
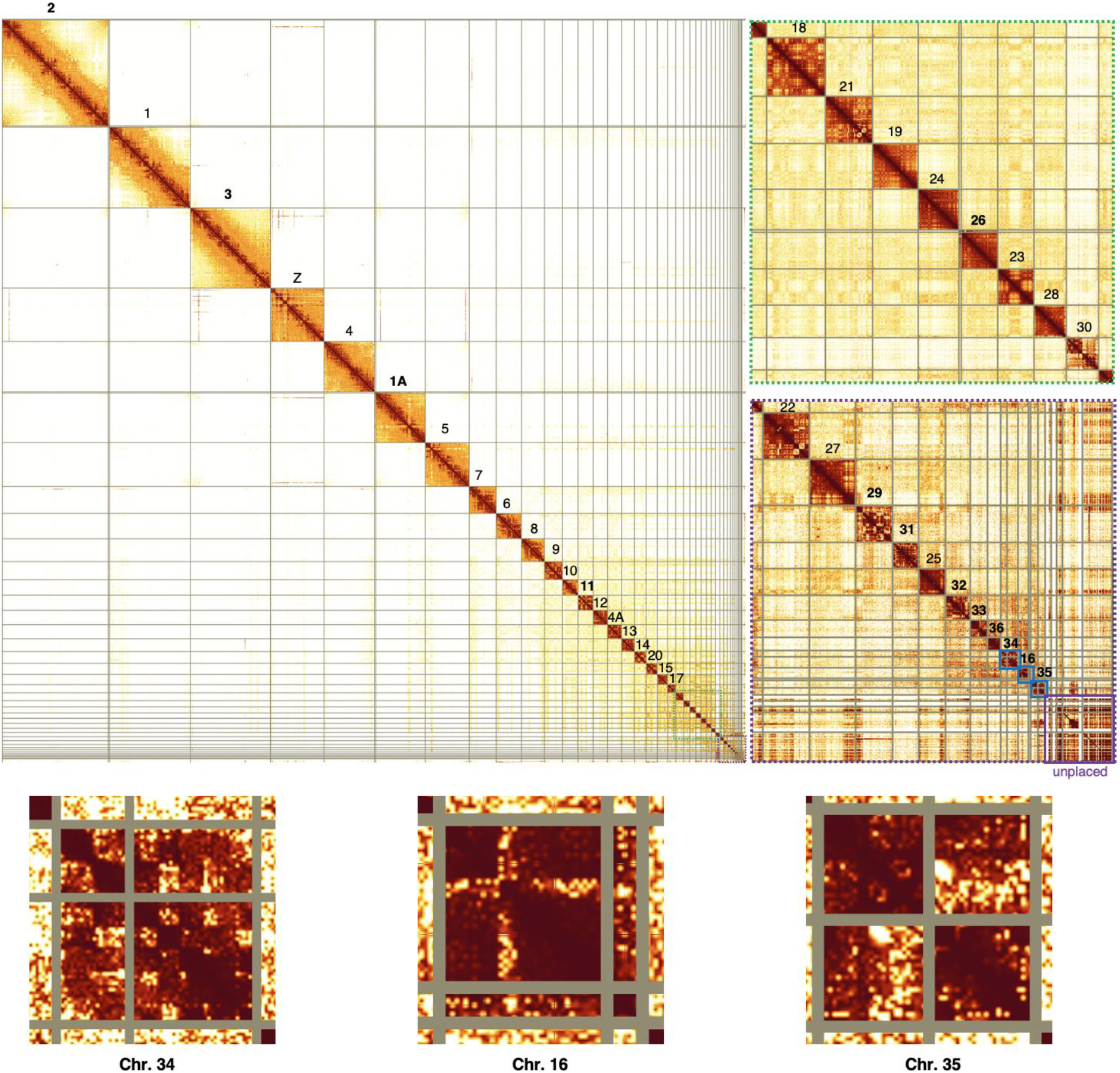
Hi-C interaction heatmap of the VGP zebra finch reassembly. Hi-C plot using PretextView of the updated bTaeGut1 v1.0 GCA_003957565.3, with newly identified chromosomal segments or additional microchromosomes named in this study highlighted in bold.

**Extended Data Fig. 3.**
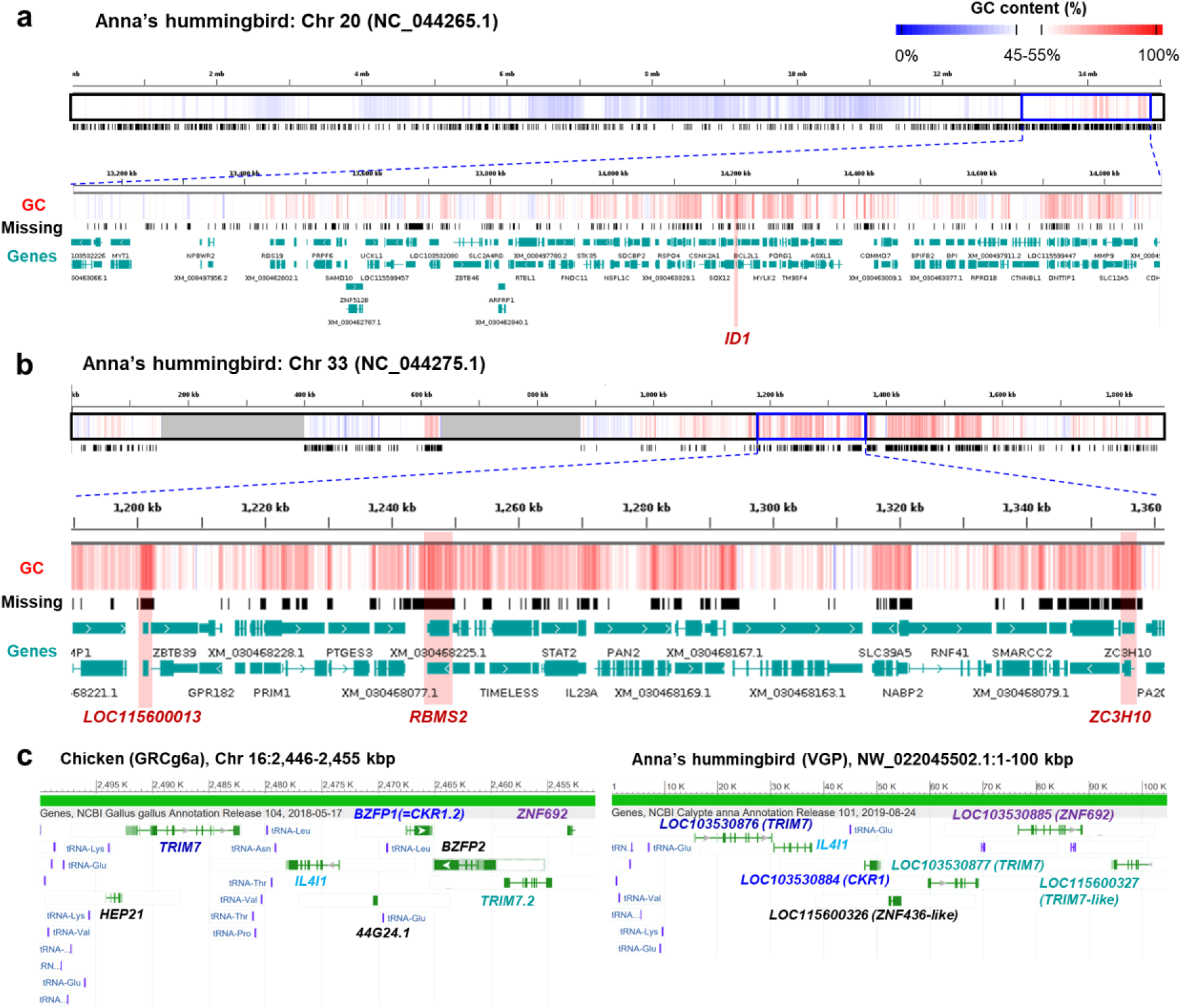
Example of missing genomic regions in Anna’s hummingbird assembly. **a,b**, Examples of previously missing GC-rich segments in Anna’s hummingbird chromosomes 20 and 33. Several genes were previously missing, either partially or completely (Red box). **c**, Fragmented chromosome 16 of VGP Anna’s hummingbird. Compared to chicken chromosome 16, one unplaced scaffold of the Anna’s hummingbird showed similar synteny, which suggests that this scaffold may be a fragment of chromosome 16 of Anna’s hummingbird, which was not assigned in the VGP assembly.

**Extended Data Fig. 4.**
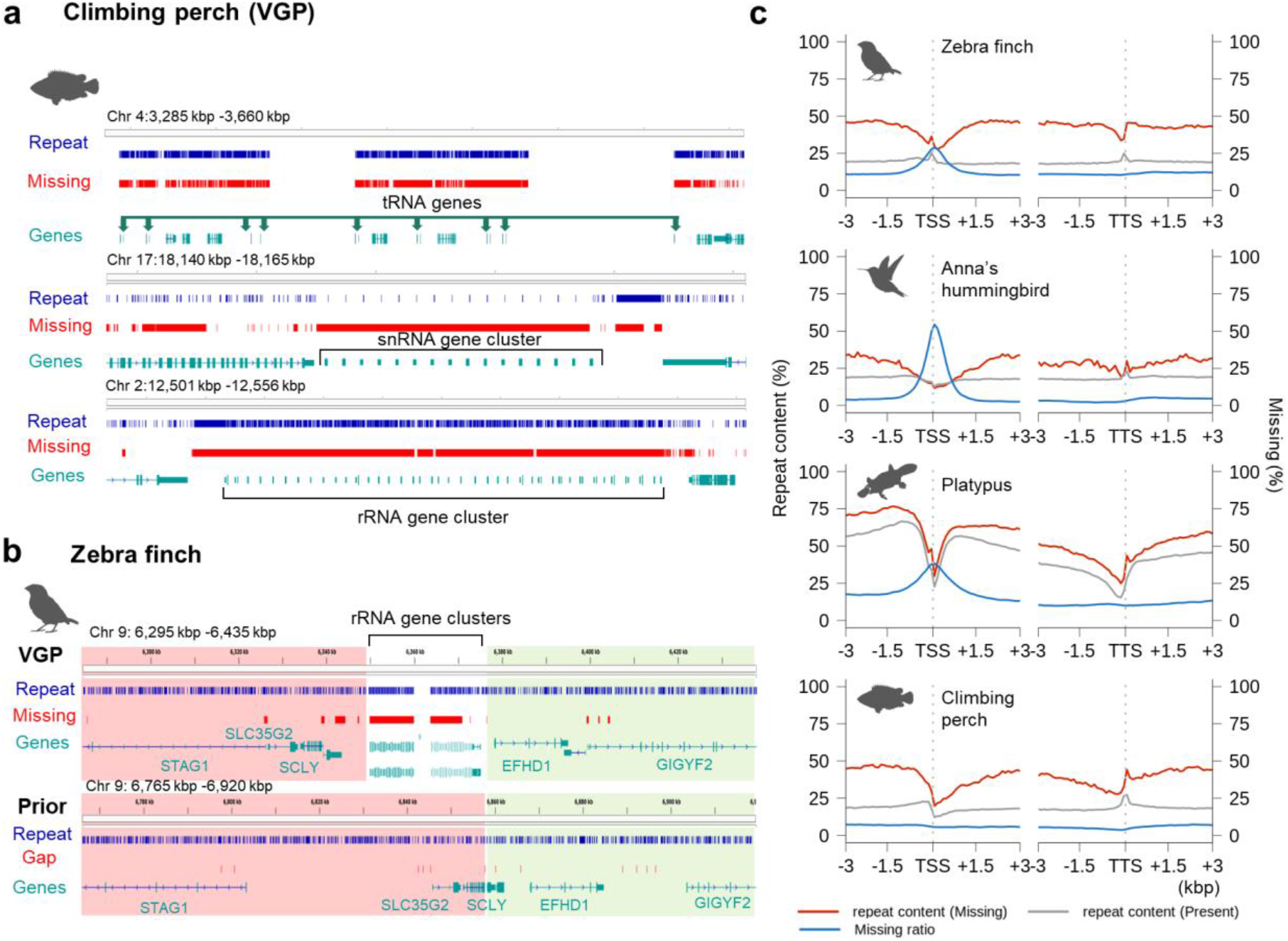
Repeat profile of previously missing genomic regions. **a,b**, Missing non-coding genes in the VGP climbing perch and zebra finch assemblies. The missing genes were within highly repetitive regions or organized repeatedly. **c**, Repeat content and missing ratio fluctuation around TSS and TTS. Red lines, repeat content of the missing blocks (90% or more missing sequences); grey lines, repeat content of the present blocks (less than 90% of missing sequences); and blue lines, the missing ratio between previous and VGP assemblies.

**Extended Data Fig. 5.**
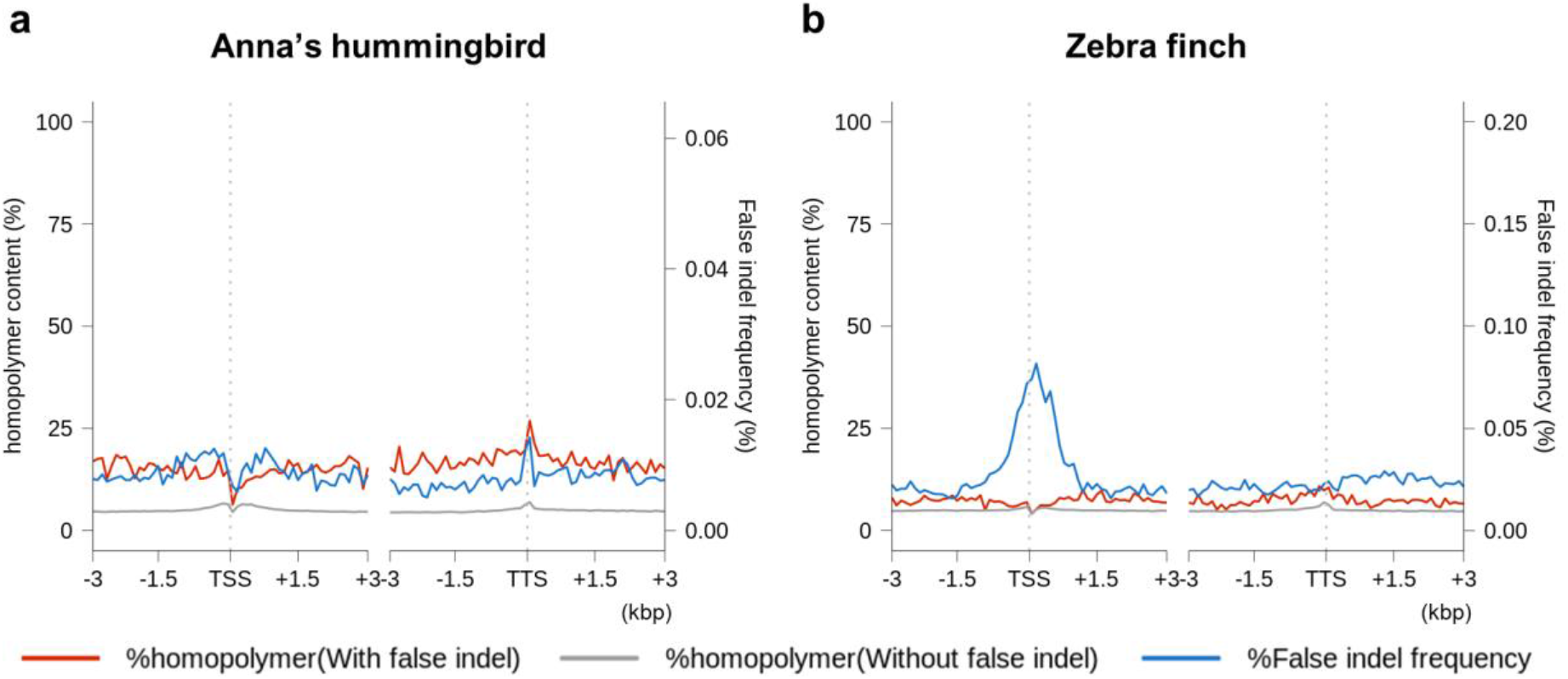
Homopolymer content and frequency of false indels near TSS and TTS. **a,b**, 100 bp consecutive blocks were extracted from the upstream and downstream 3 kbp regions of genes and classified depending on the presence of false indels. Red lines, homopolymer content of the blocks with false indels. Grey lines, blocks without false indels. Blue line, frequency of false indels.

**Extended Data Fig. 6.**
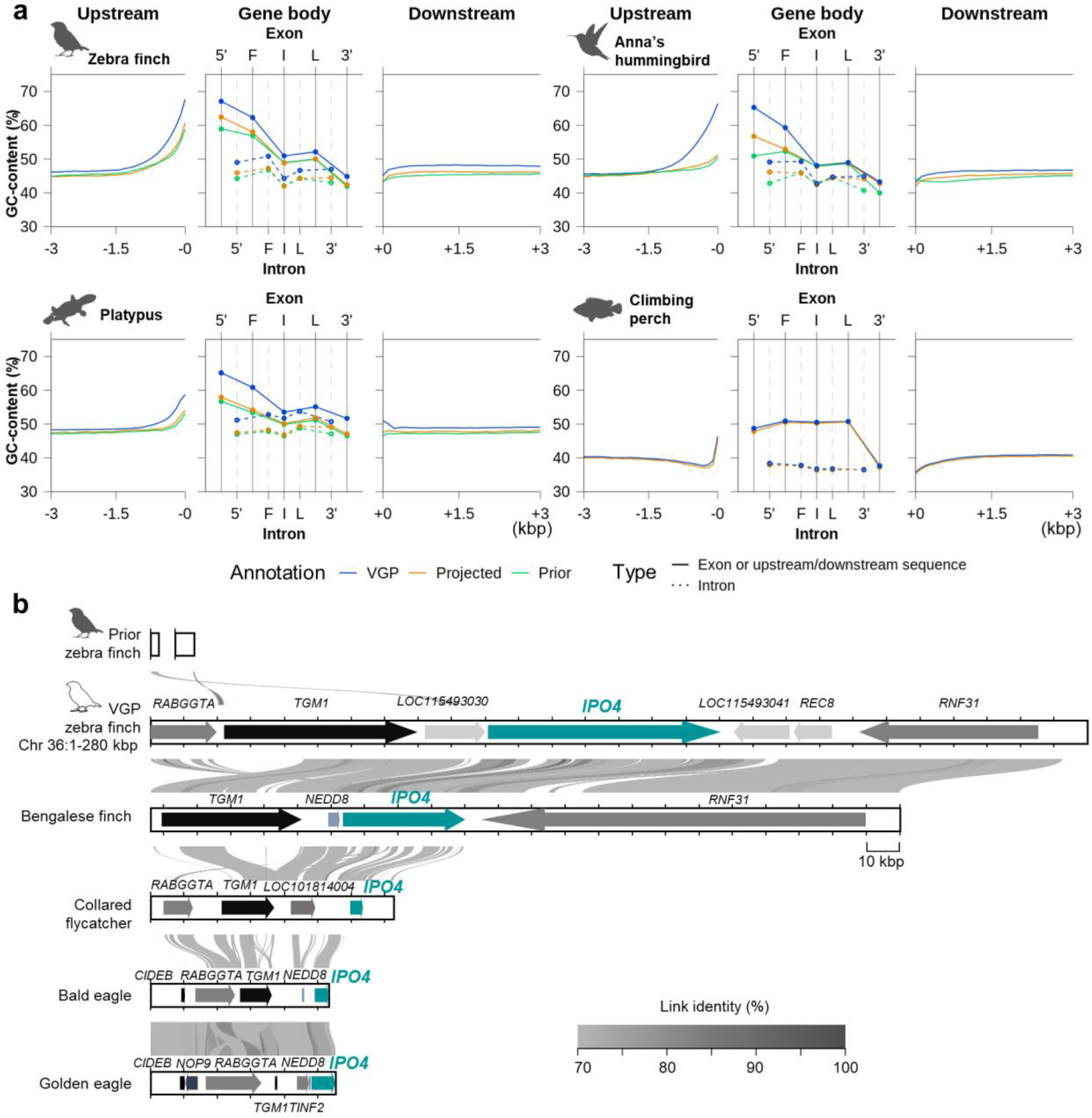
Improvements of VGP annotations compared to prior annotations. **a**, Average GC-content of protein coding genes from VGP (blue), projected (yellow), and prior (green) annotations. 5’: 5’UTR, F: First coding, I: Internal coding, L: last coding, 3’: 3’UTR exon or intron. **b**, Alignment of genomic regions including the *IPO4* gene in several bird assemblies. Visualization of conserved synteny was based on AliTV^72^.

**Extended Data Fig. 7.**
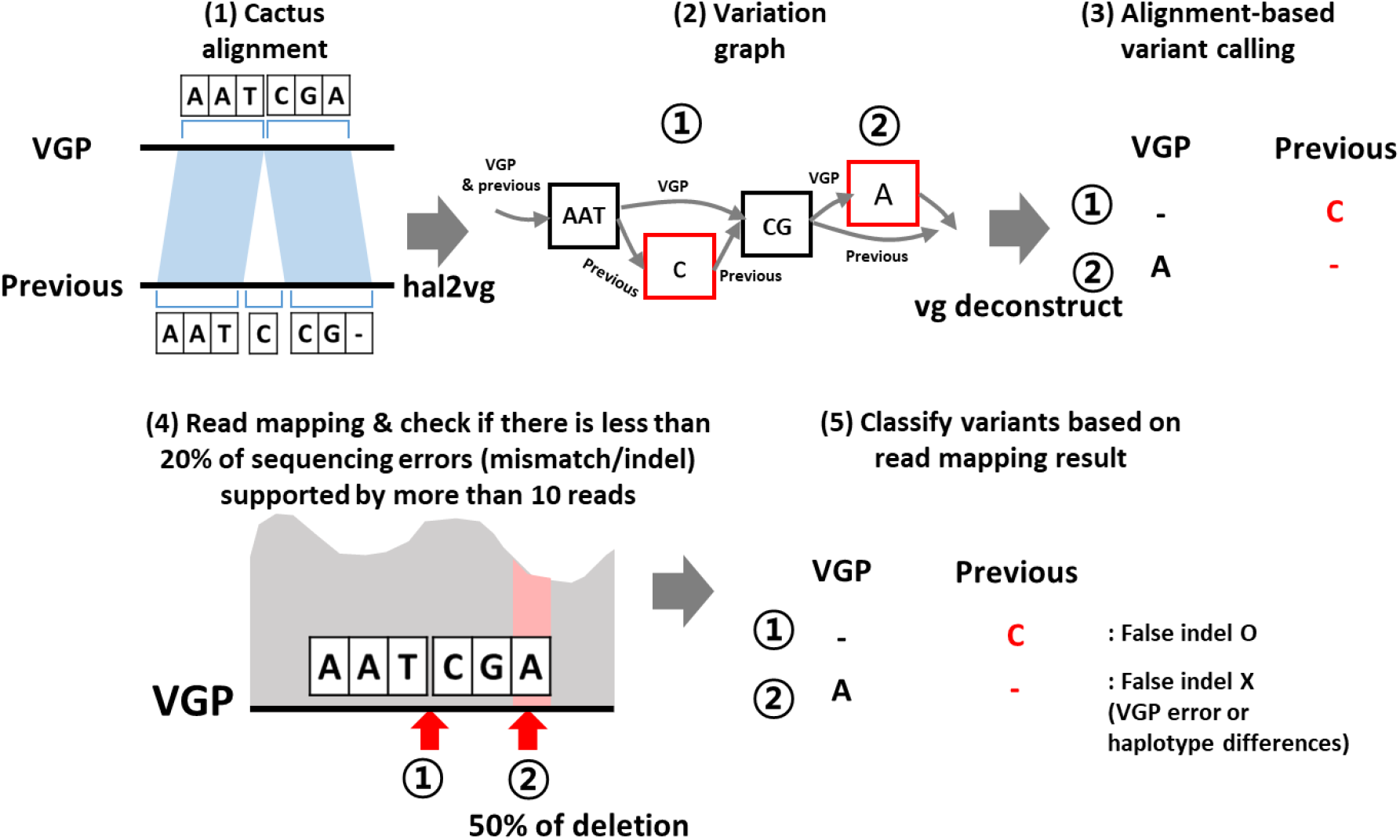
Detection of false indel (and false SNP) from cactus alignment and mpileup result. We transformed the cactus alignment into a variation graph using hal2vg. Next, variants based on genome-wide alignment were called with the deconstruct of vg toolkit using VGP primary assembly as a reference. The genomic coordinates of potential VGP assembly errors or heterozygous alleles were collected from the mpileup results with a threshold 20% and +/−2 bp flanking sequences. The variants called from genome-wide alignment were excluded when their size was more than 10 bp or they overlapped the genomic coordinates of potential VGP assembly errors or heterozygous alleles.

**Extended Data Fig. 8.**
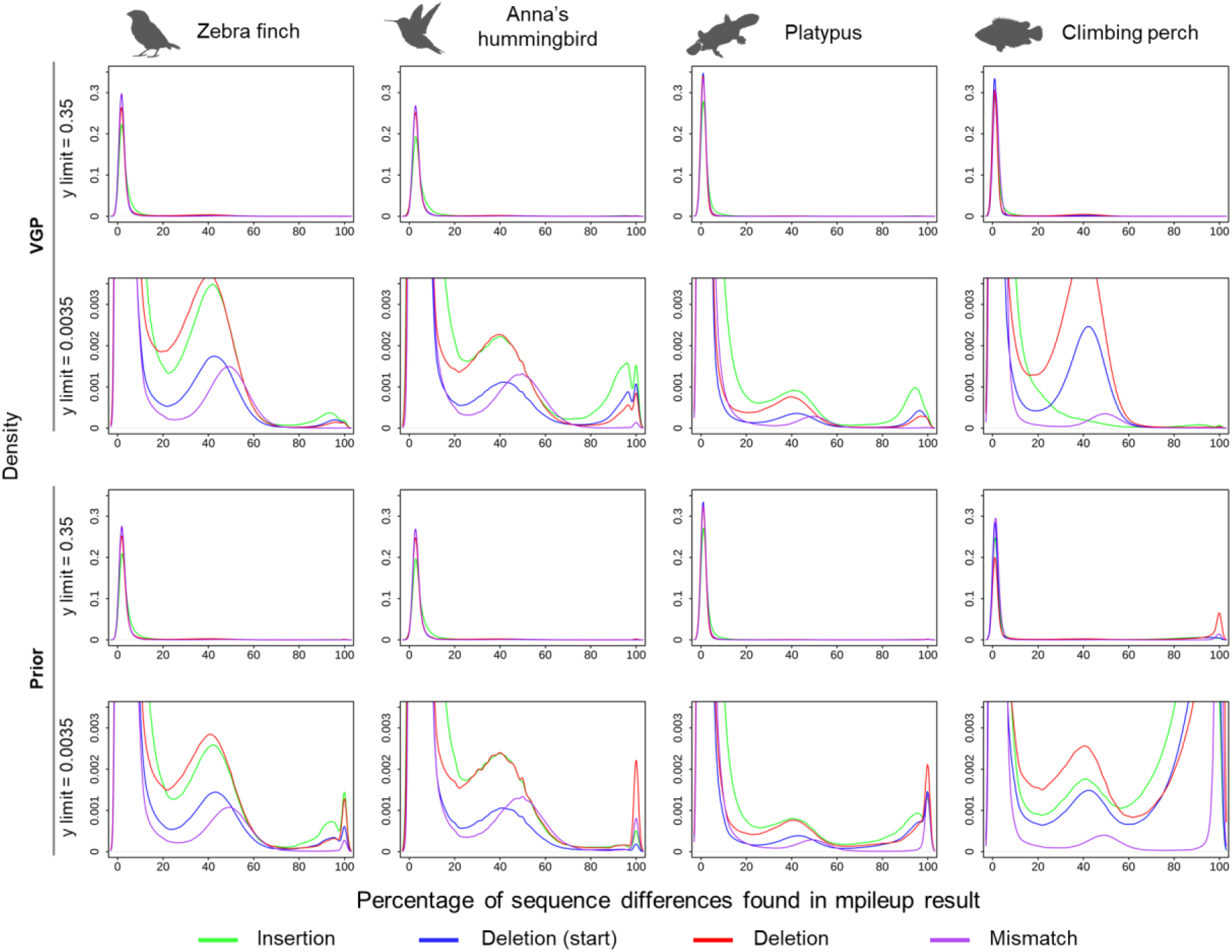
Density plots of the proportion of sequence differences found by mpileup results of 10x Genomics linked-read libraries mapped to the VGP primary and prior assemblies. From genome-wide mpileup results, loci with 10 or more reads and one or more sequence differences were collected. Proportion of sequence differences was calculated by the number of sequence differences divided by the number of reads mapped on each locus. In the cases of deletions, the read bases right before the deletion (blue) and the following deleted (red) were counted separately. From the result, we concluded that a 20% and an 80% threshold are suitable for collecting heterozygous alleles or potential VGP assembly errors, respectively.

**Extended Data Fig. 9.**
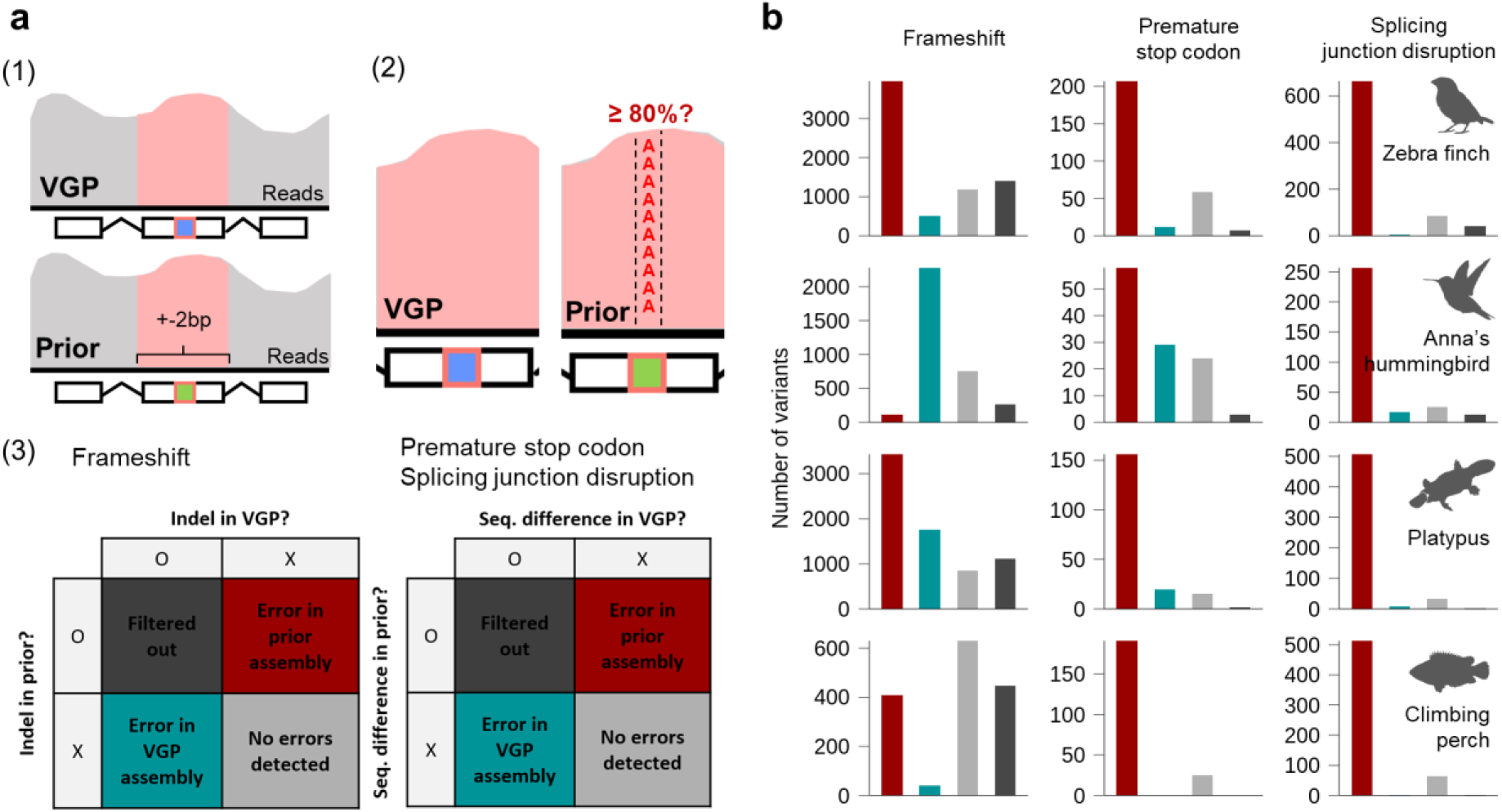
Detection of sequence-level false gene losses. **a**, Summary of the method to detect sequence-level fasle gene losses. Based on the difference between the VGP and previous assemblies, SAMTools’ mpileup was used to further classify these false losses into four categories. Error in prior assembly (FGL, red): a sequencing error or indel found in the prior assembly, Error in VGP assembly (blue): a sequencing error or indel found in the VGP assembly. No errors detected (light grey): both assemblies did not show a sequencing error or indel supported by read mapping data. Filtered out (dark grey): both VGP and previous assemblies showed sequencing errors. **b**, Number of frameshift, splicing junction disruption, and premature stop codon errors.

